# Modelling Predictive Coding in the Primary Visual Cortex (V1): Layer 4 Receptive Field Properties in a Balanced Recurrent Spiking Neuronal Network

**DOI:** 10.1101/2025.10.20.683584

**Authors:** Elnaz Nemati, Catherine Davey, Hamish Meffin, Anthony N. Burkitt

**Affiliations:** Department of Biomedical Engineering, The University of Melbourne, Victoria, Australia; Graeme Clark Institute, The University of Melbourne, Victoria, Australia

## Abstract

Understanding how the cortex encodes sensory input in a biologically efficient and computationally robust manner remains a central question in neuroscience. Predictive coding offers a compelling theoretical framework for such cortical processing, but existing models lack the biological detail to fully explain the function of the cortical microcircuits. This study introduces a spiking neural network model of layer 4 of the primary visual cortex (V1), grounded in predictive coding principles, to clarify how the thalamorecipient layer transforms feedforward input into prediction-error-like signals under realistic excitatory–inhibitory constraints and to yield testable circuit-level predictions. The model integrates structured feedforward input, distinct excitatory and inhibitory populations, and balanced lateral connectivity to simulate spontaneous and stimulus-driven activity. Network responses are systematically examined under spatially unstructured noise input and structured grating stimuli. Neural membrane potentials encode real-time reconstruction errors between external input and internal estimates, with spikes dynamically correcting these mismatches. The network reproduces hallmark in vivo features, including irregular spontaneous activity, sparse and selective responses, and emergent orientation and phase tuning. Excitatory-Inhibitory (E-I) balance was maintained across conditions, with inhibitory neurons exhibiting tighter input coupling than excitatory neurons. Furthermore, the network exhibited contrast-dependent modulation of firing rates and E-I balance, dynamically adjusting its activity to changes in input strength. Decoding analyses demonstrates that structured inputs can be robustly reconstructed under moderate noise levels, although decoding fidelity declines sharply under severe corruption. Together, these results suggest that cortical layer 4 may serve as a structured sensory encoding stage in a hierarchical predictive coding system, providing a biologically grounded foundation for modeling prediction error computations in higher cortical areas.

**Author summary:** In this Study, we present a biologically grounded spiking neural network model of layer 4 of the primary visual cortex, built within the predictive coding framework. The aim is to better understand how this early cortical layer encodes sensory information while maintaining realistic neural dynamics. Many predictive coding models focus on higher cortical layers 2/3 and overlook layer 4’s role. To address this, we develop here a network that integrates structured feedforward input via Gabor-filtered receptive fields, distinct excitatory and inhibitory populations, and fixed lateral connectivity, all adhering to Dale’s law. The model reproduces several *in vivo* features observed in layer 4 of the visual cortex, including sparse, irregular spiking, emergent orientation and phase tuning, and contrast-dependent firing. Notably, excitation and inhibition are dynamically balanced across input conditions without requiring synaptic learning. We also show that decoding performance remains robust under moderate noise levels, supporting that layer 4 provides a stable sensory foundation for higher-level prediction. This model offers a biologically realistic implementation of prediction error computation and sets the stage for hierarchical extensions that include feedback and learning. Overall, this work provides insights into how structured sensory representations and balance emerge in cortical microcircuits through architecture alone.

## 1 Introduction

Understanding sensory processing in the brain requires models that can efficiently extract relevant features from complex inputs while maintaining biological realism. Predictive coding provides an influential theoretical framework by proposing that the brain continuously generates predictions about sensory input and updates neuronal activity to minimize the mismatch between expectation and sensory input [1–3]. In this framework, neural responses primarily reflect prediction errors, i.e., differences between expected and observed input, rather than a direct encoding of raw sensory signals. This principle supports efficient information transmission and robust inference under uncertainty. Importantly, testing the biological plausibility of predictive coding requires identifying how specific cortical layers implement these computations.

In cortical hierarchies, layer 4 is widely recognized as the principal recipient of feedforward sensory input, serving as a critical gateway for constructing early representations of sensory features [15, 16, 18]. Within predictive coding frameworks, this role positions layer 4 as the entry point for sensory evidence, which is then passed on to layer 2/3 where it can be integrated with top-down predictions to guide perception.

However, many predictive coding models have paid little attention to this layer, either omitting it entirely or treating it as a passive relay, while focusing instead on computations in layers 2/3 and layer 5. Canonical predictive coding architectures, such as those proposed by Bastos et al. [17], reinforce this view by simplifying layer 4 to a feedforward pathway from the Lateral Geniculate Nucleus (LGN) to layer 2/3. As a consequence, the specific computations performed within layer 4 and their contribution to predictive coding remain poorly understood. In particular, it is still unclear how this layer combines incoming sensory evidence with local lateral dynamics to form reliable representations for comparison with descending predictions.

Given its central role in sensory processing, a large number of spiking neuron models have sought to capture the computations of layer 4. These models highlight features such as push-pull mechanisms [19], orientation selectivity [20], sparse spiking [3], and natural scene encoding [21], demonstrating the importance of this layer for understanding cortical input processing. However, while these studies provide valuable insight into the physiology of layer 4, they generally do not consider its function within a predictive coding framework. Table 1 summarizes these models and the biological features they implement, such as synaptic plasticity, LGN modeling, and predictive behavior.

**Table 1.**
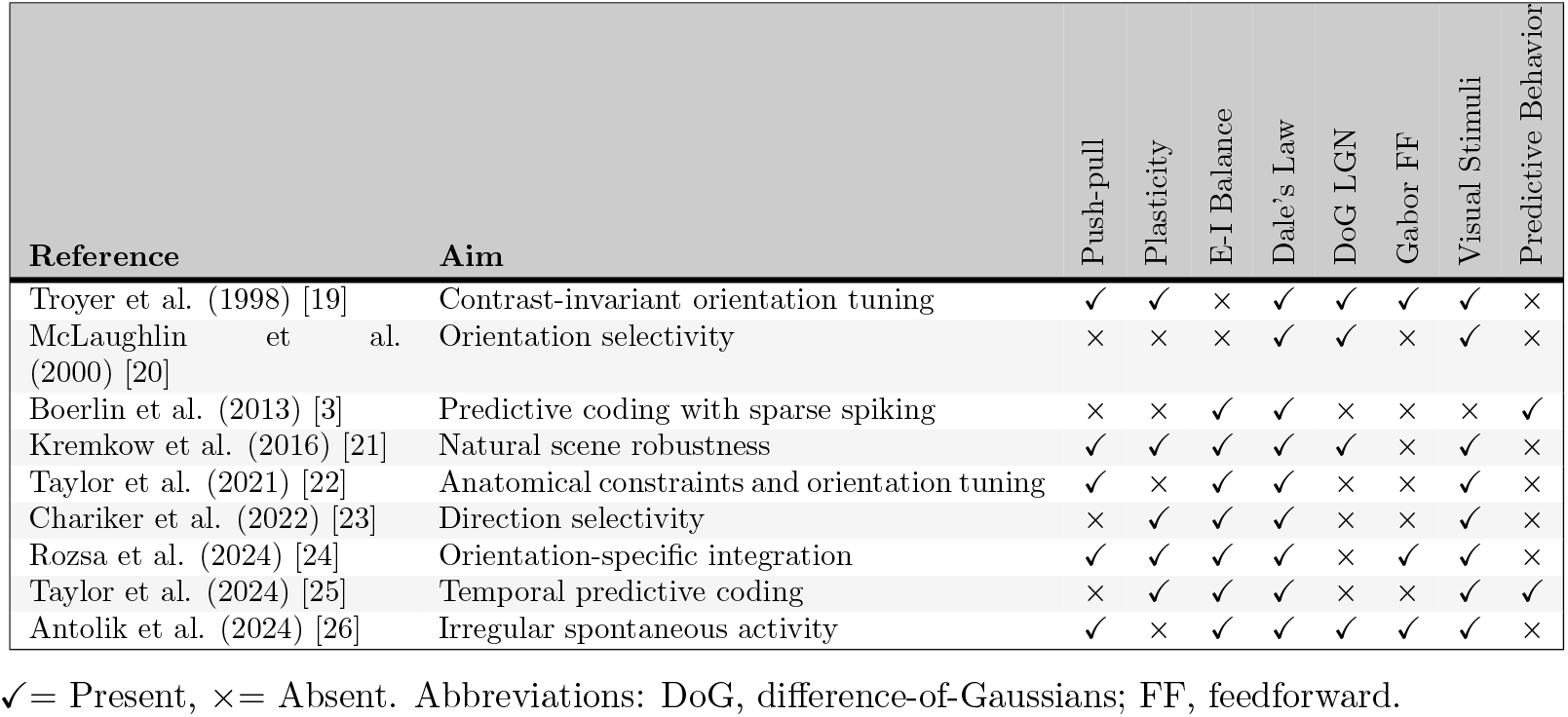
Summary of Layer 4 Spiking Neuron Models.

| Reference | Aim | Push-pull | Plasticity | E-I Balance | Dale's Law | DoG LGN | Gabor FF | Visual Stimuli | Predictive Behavior |
| --- | --- | --- | --- | --- | --- | --- | --- | --- | --- |
| Troyer et al. (1998) [19] | Contrast-invariant orientation tuning | ✓ | ✓ | × | ✓ | ✓ | ✓ | ✓ | × |
| McLaughlin et al. (2000) [20] | Orientation selectivity | × | × | × | ✓ | ✓ | × | ✓ | × |
| Boerlin et al. (2013) [3] | Predictive coding with sparse spiking | × | × | ✓ | ✓ | × | × | × | ✓ |
| Kremkow et al. (2016) [21] | Natural scene robustness | ✓ | ✓ | ✓ | ✓ | ✓ | × | ✓ | × |
| Taylor et al. (2021) [22] | Anatomical constraints and orientation tuning | ✓ | × | ✓ | ✓ | × | × | ✓ | × |
| Chariker et al. (2022) [23] | Direction selectivity | × | ✓ | ✓ | ✓ | × | × | ✓ | × |
| Rozsa et al. (2024) [24] | Orientation-specific integration | ✓ | ✓ | ✓ | ✓ | × | ✓ | ✓ | × |
| Taylor et al. (2024) [25] | Temporal predictive coding | × | ✓ | ✓ | ✓ | × | × | ✓ | ✓ |
| Antolik et al. (2024) [26] | Irregular spontaneous activity | ✓ | × | ✓ | ✓ | ✓ | ✓ | ✓ | × |

✓ = Present, × = Absent. Abbreviations: DoG, difference-of-Gaussians; FF, feedforward.

These models, summarized in Table 1, have advanced our understanding of the neural computations carried out in layer 4, but each has typically focused on isolated features such as orientation selectivity, push–pull mechanisms, or sparse spiking. Importantly, few of these approaches consider predictive neural dynamics directly or incorporate biologically realistic decoding weights. This limitation motivates turning to the broader class of predictive coding models in spiking networks, where related challenges of robustness, dynamics, and biological realism have been investigated.

The spike-based predictive coding framework introduced by Boerlin et al. [3] has inspired numerous extensions that aim to overcome its computational limitations and improve biological plausibility. These studies have explored robustness, synaptic delays, spontaneous activity, learning, and population heterogeneity, but few have combined these features with structured sensory input, receptive field alignment, or layer-specific architecture. A summary of these extensions is provided in Table 2, grouped according to the particular features that characterize them.

**Table 2.** Summary of variants of predictive coding and their biological features.

| Reference | Focus | Dale's Law | Robustness | Learning | Natural Images | Gabor receptive fields | Orientation/Phase Tuning |
| --- | --- | --- | --- | --- | --- | --- | --- |
| **Robustness and Compensation** |  |  |  |  |  |  |  |
| Fiquet et al. (2016) [27] | Robustness in spiking networks | × | ✓ | × | × | × | × |
| Barrett et al. (2016) [28] | Adaptation to neuron loss | × | ✓ | × | ✓ | ✓ | ✓ |
| Gutierrez et al. (2019) [36] | Population adaptation in balanced networks | × | ✓ | × | × | × | × |
| Zeldenrust et al. (2021) [29] | Heterogeneity improves coding | × | ✓ | × | × | × | × |
| **Temporal Dynamics and Spontaneous Activity** |  |  |  |  |  |  |  |
| Chalk et al. (2016) [30] | Oscillations from synaptic delays | ✓ | ✓ | × | × | × | × |
| Koren and Denève (2017) [32] | Spontaneous activity as prediction | × | × | × | × | × | × |
| Recanatesi et al. (2021) [31] | Predictive learning in low-dimensional space | × | ✓ | ✓ | × | × | × |
| **Learning and Plasticity** |  |  |  |  |  |  |  |
| Alemi et al. (2017) [33] | Learning dynamic transformations | × | × | ✓ | × | × | × |
| Kushnir and Denève (2019) [34] | Learning temporal structure in spiking networks | × | × | ✓ | × | × | × |
| Brendel et al. (2020) [35] | Spike-by-spike predictive learning | × | × | ✓ | × | × | × |
| **Biological Realism** |  |  |  |  |  |  |  |
| Chalk et al. (2017) [37] | Receptive field adaptation to sensory noise | × | × | × | × | ✓ | ✓ |
| Mikulasch et al. (2023) [38] | Dendritic predictive computation | × | × | × | × | × | × |
| This study | Layer 4 predictive coding with natural input | ✓ | ✓ | × | ✓ | ✓ | ✓ |

✓ = Present, × = Absent.

As Table 2 highlights, robustness has been a central focus. Fiquet et al. [27] showed that predictive coding networks can tolerate neuron loss and synaptic noise through redundancy in recurrent dynamics. Similarly, Barrett et al. [28] demonstrated that tightly balanced networks can reassign functional roles in response to perturbations. Heterogeneity, as explored by Zeldenrust et al. [29], improves both stability and coding precision across conditions.

The spike-based predictive coding framework introduced by Boerlin et al. [3] has inspired numerous extensions addressing its computational limitations and efforts to improve biological realism. These studies have explored robustness, synaptic delays, spontaneous activity, learning, and population heterogeneity, but few have combined these features with structured sensory input, receptive field alignment, or layer-specific architecture.

Robustness has been a central focus. Fiquet et al. [27] showed that predictive coding networks can tolerate neuron loss and synaptic noise through redundancy in recurrent dynamics. Similarly, Barrett et al. [28] demonstrated that tightly balanced networks can reassign functional roles in response to perturbations. Heterogeneity, as explored by Zeldenrust et al. [29], improves both stability and coding precision across conditions.

Several studies have addressed temporal processing. Chalk et al. [30] introduced synaptic delays, showing that oscillations can naturally arise from predictive dynamics. Recanatesi et al. [31] extended predictive learning to uncover low-dimensional latent representations in spiking networks. In addition, Koren and Denève [32] proposed that spontaneous activity reflects internal model dynamics during idle states.

A number of studies have extended predictive coding models by incorporating neural learning. Alemi et al. [33] proposed local plasticity rules for learning dynamic transformations, while Kushnir and Denève [34] showed that recurrent spiking networks can learn temporal input structure. Brendel et al. [35] introduced a spike-by-spike predictive learning rule, and Gutierrez and Denève [36] investigated adaptation in balanced networks. These studies highlight the importance of learning for predictive coding frameworks; however, as our focus here is on the biophysical implementation of predictive dynamics in layer 4, the question of learning lies outside the scope of this work.

Chalk et al. [37] addressed adaptation to noise in receptive fields, and Mikulasch et al. [38] explored how dendritic integration supports predictive computation. Despite these advances, biological realism remains a challenge across much of this work: many models are evaluated only on simplified stimuli rather than naturalistic image inputs, and they often omit key constraints such as Dale’s Law, tuning diversity, or responsiveness to natural scenes.

Taken together, existing spiking models of layer 4 and extensions of predictive coding have each provided valuable insights, but they typically address only isolated aspects of the problem. For example, classical models of layer 4 such as Troyer et al. [19], Kremkow et al. [21], and Rozsa et al. [24] emphasize contrast invariance, orientation tuning, and natural scene robustness. Yet, they neglect predictive dynamics and do not enforce biologically realistic constraints such as balanced excitation–inhibition. Conversely, predictive coding frameworks including Boerlin et al. [3], Chalk et al. [30], and Kushnir and Denève [34] have explored robustness, oscillations from synaptic delays, and temporal learning, but these are typically evaluated on highly simplified inputs and rarely incorporate structured receptive fields aligned with V1 physiology. Even more biologically detailed approaches, such as Chalk et al. [37] or Mikulasch et al. [38], stop short of combining naturalistic image-driven input with predictive coding dynamics. As a result, it remains unclear how predictive coding can be instantiated in a biologically realistic layer 4 microcircuit that processes naturalistic sensory inputs.

To address this gap, we introduce a spiking neural network model of cortical layer 4 in the primary visual cortex that builds on the predictive coding framework of Boerlin et al. [3]. The model encodes reconstruction errors through spiking dynamics while incorporating biologically grounded features often omitted in prior work. Specifically, it enforces Dale’s Law and excitatory–inhibitory balance and structures the feedforward drive using Gabor-filtered input. Gabor functions provide a well-established description of V1 receptive fields [5–9], capturing orientation and spatial frequency selectivity, hallmark properties of layer 4 simple cells in primary visual cortex. This ensures that natural image processing is modeled in a way that reflects the functional architecture of V1. Moreover, fixed lateral connectivity based on decoding weights gives rise to emergent balance, sparsity, and feature selectivity without requiring synaptic learning, thereby providing a biologically plausible instantiation of predictive coding in a canonical cortical circuit.

In the sections that follow, the model is presented in detail, evaluated for its computational properties under naturalistic input, and compared with both existing predictive coding frameworks and layer-specific spiking models, highlighting the novel contributions of our approach.

## 2 Methods

### 2.1 Network Structure

The network comprises distinct excitatory and inhibitory neuron populations that adhere to Dale’s law and follow a canonical 4:1 excitatory-to-inhibitory ratio, reflecting the cellular composition observed in neocortical microcircuits [10, 11]. All synaptic connections are confined laterally within layer 4 to model local recurrent interactions, while top-down and feedback pathways are excluded, as these predominantly target supragranular and infragranular layers rather than layer 4 [16, 17].

Each neuron receives input from simulated visual stimuli through small image patches, such as gratings or natural scenes. The input is decomposed into ON and OFF pathways, reflecting the well-established segregation of positive and negative luminance signals in the retina and lateral geniculate nucleus (LGN) [12–14]. For each image patch, the mean luminance is treated as the local background. Pixels brighter than this background drive the ON pathway, while pixels darker than the background drive the OFF pathway, corresponding to positive and negative luminance deviations, respectively. These ON and OFF contrast signals are converted into Poisson spike trains and provide the LGN-like afferent input to the layer 4 cortical network.

All excitatory layer-4 neurons receive input from both ON and OFF pathways, weighted according to the structure of their Gabor receptive fields. These receptive fields are parameterized by their five features: orientation, phase, spatial frequency, size, and position. The ON and OFF components of each Gabor filter separately determine the synaptic weights from ON and OFF LGN channels.

The overall network architecture is illustrated in Fig 1. Panel (A) shows how the Gabor structure connects ON and OFF LGN pathways to excitatory neurons. Each receptive field acts as a decoding filter integrating input from the LGN. Neurons with similar receptive field structures are more strongly connected, forming spatial clusters (highlighted in black). In contrast, connections between dissimilar receptive field groups are weaker, indicated by dashed lines.

**Fig 1.**
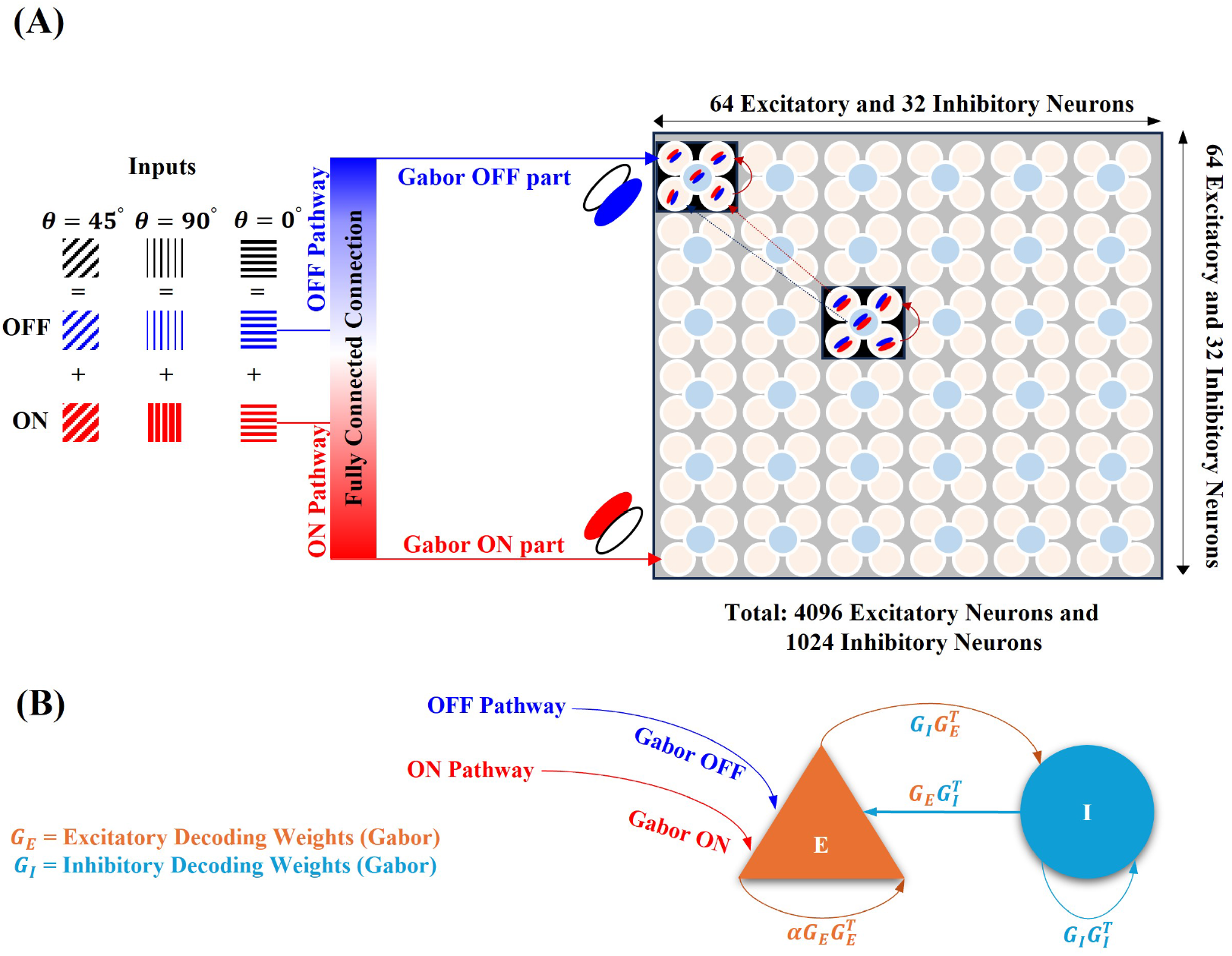
Overview of the layer 4 predictive coding network architecture. **(A)** Input stimuli (e.g., gratings or natural scenes) are decomposed into ON (red) and OFF (blue) LGN pathways, each projecting to excitatory cortical neurons through Gabor-filtered decoding weights (parameter definitions given in Table 3). Each image patch is split into ON and OFF components that are *fully connected* to all excitatory cortical neurons; however, connection strength is scaled by the similar- ity of their Gabor features. Neurons with similar orientation and phase form strongly connected local sub-networks (black boxes), whereas neurons with dissimilar features have weak or negligible connections (dashed lines).The two highlighted boxes illustrate sub-networks tuned to similar orientations but opposite (reversed) spatial phases, consistent with feature-dependent lateral connectivity. Lateral connectivity between excitatory (E, orange) and inhibitory (I, sky blue) populations follows an all-to-all structure, with synaptic weights determined by their respective Gabor decoding weights. Excitatory and inhibitory weight matrices are denoted as *G*_E_ and *G*_I_, respectively, and connection strengths are modulated by feature similarity to maintain balanced excitation and inhibition across the layer.

Panel (B) depicts the lateral interactions within the excitatory and inhibitory populations. Excitatory neurons (red) and inhibitory neurons (blue) interact through structured connectivity defined by their Gabor decoding weights. These weight matrices govern both E-to-I and I-to-E coupling, as well as E-to-E and I-to-I lateral inhibition.

**Table 3.** Gabor filter parameters: For a 16 × 16 pixel receptive field with 16 pixels per degree.

| Parameter | Description |
| --- | --- |
| Orientation $\theta$ | Randomly sampled from $0^\circ$ to $180^\circ$ to span the full range of orientation preferences. |
| Phase $\psi$ | Discretely chosen from $0^\circ$, $90^\circ$, $180^\circ$, or $270^\circ$ to support phase-specific encoding. |
| Spatial frequency $\lambda$ | Set to 0.8 cycles per degree, corresponding to a wavelength of approximately 19.6 pixels. |
| Envelope width $\sigma_x$ and $\sigma_y$ | Fixed at $0.17^\circ$ of visual field, equivalent to approximately 2.67 pixels. |
| Aspect ratio $\gamma$ | Uniformly sampled from 1.7 to 5, reflecting variability in biological receptive field elongation. |

**Table 4.** Model parameters used for simulations of a V1 network.

| Parameter | Description and Value |
| --- | --- |
| Input patch size | $16 \times 16$ pixels |
| Number of LGN neurons ($N_{\text{LGN,ON}}, N_{\text{LGN,OFF}}$) | 256 ON, 256 OFF |
| Number of excitatory neurons ($N_E$) | 4096 |
| Number of inhibitory neurons ($N_I$) | 1024 |
| Simulation time step ($\Delta t$) | 0.1 ms |
| Membrane time constant ($\tau_m$) | 10 ms (exc), 10 ms (inh) |
| Leak potential ($E_L^{(E)}, E_L^{(I)}$) | -80 mV (exc), -80 mV (inh) |
| Membrane resistance ($R_m^{(E)}, R_m^{(I)}$) | 100 MΩ (exc), 50 MΩ (inh) |
| Synaptic time constants ($\tau_E, \tau_I$) | 5 ms (exc), 10 ms (inh) |
| Time constant for rate decoder ($\tau_d$) | 10 ms |
| Reset potential ($V_r^{(E)}, V_r^{(I)}$) | -80 mV (exc), -80 mV (inh) |
| Refractory period ($\tau_{r,E}, \tau_{r,I}$) | 2 ms (exc), 0.5 ms (inh) |
| Gaussian noise term ($n_\sigma$) | Zero mean, small variance |

This model allows structured, orientation- and phase-selective processing of input stimuli within a balanced recurrent network. The following sections provide a detailed description of the input pre-processing pipeline, decoding weight construction, neuronal and synaptic dynamics, and stimulus presentation protocol.

### 2.2 Layer-4 Input Pre-processing and Decoding Weights

To simulate early visual input to cortical layer 4, the network receives image patches from either natural scenes or synthetic stimuli such as oriented gratings. These inputs are preprocessed using biologically inspired transformations to approximate retinal and LGN processing, as illustrated in Fig. 2.

**Fig 2.**
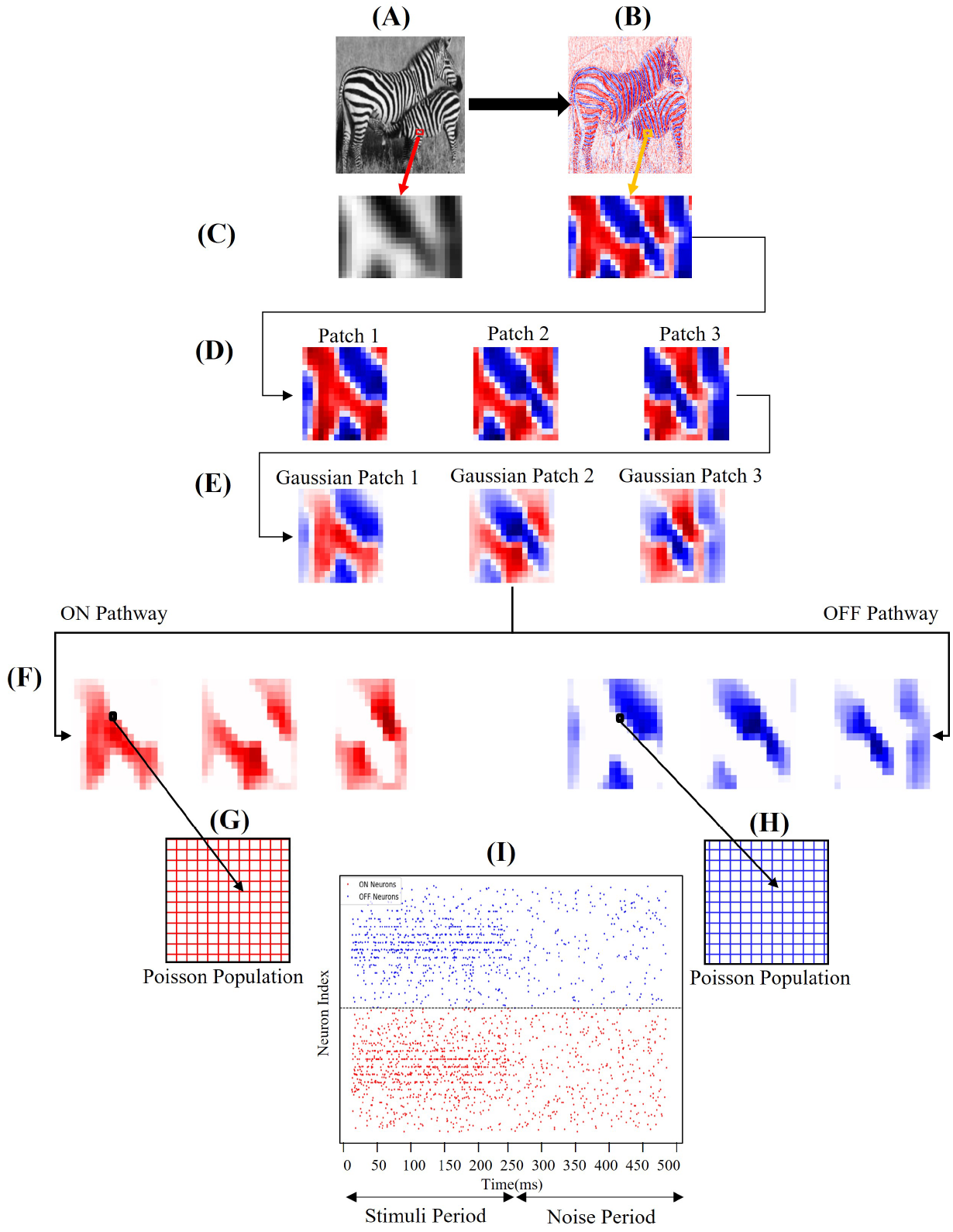
Pre-processing pipeline for input stimuli. **(A)** Example grayscale natural image (zebra) used as model input. **(B)** Images are whitened following Olshausen and Field [39] to remove second-order correlations and normalize contrast. **(C)** A 16 × 26 region of interest is extracted from the image for subsequent processing. **(D)** This region is subdivided into overlapping 16 × 16 patches that simulate localized receptive-field sampling across the visual field. **(E)** Each patch is convolved with a Gaussian filter to approximate spatial integration in the LGN and smooth luminance variations [1]. **(F)** The filtered signal is decomposed into ON (red) and OFF (blue) channels according to pixel polarity, corresponding to positive and negative luminance deviations from the local mean. **(G–H)** Each ON and OFF pixel is mapped onto a corresponding neuron within separate Poisson populations, converting luminance intensity into spike trains. The ON and OFF populations are arranged on separate 16 × 16 neural grids to represent parallel input pathways. **(I)** Raster plot showing ON (red) and OFF (blue) spikes over a 500 ms simulation. The first half (0–250 ms) corresponds to stimulus presentation, and the second half (250–500 ms) depicts background activity after stimulus offset.

Image whitening enhances contrast and removes pixel-level correlations, approximating early retinal and LGN processing [40–43]. In biological vision, these operations correspond to the center–surround organization of retinal ganglion cells and LGN neurons, which decorrelate natural visual input and emphasize edges, thereby increasing the efficiency of sensory encoding [40, 41]. To model this process, a zero-phase whitening filter was applied in the Fourier domain, defined by the power spectrum:

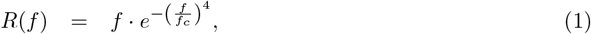

where *f*_*c*_ = 200 denotes the characteristic spatial frequency for each image. This filter removes low-frequency correlations while preserving fine-scale structure, producing a spatial response profile that closely resembles the center–surround receptive fields of LGN neurons [39, 40]. Following whitening, the images were normalized to a fixed variance (0.2) and convolved with an additional Gaussian kernel to approximate further spatial integration by LGN cells, following the approach of Rao and Ballard [1]. Thus, this preprocessing step provides a biologically realistic front-end transformation that mimics early visual encoding before cortical processing.

The resulting signal is then split into ON and OFF channels: pixels with positive intensities are routed through the ON pathway, and negative pixels through the OFF pathway. Each channel is converted into spike trains using independent Poisson processes with rates *λ*_*ON*_ and *λ*_*OFF*_, respectively. Baseline firing rates of 17 Hz (ON) and 8 Hz (OFF) were added to reflect tonic LGN activity under spontaneous conditions [44].

The spiking outputs from the ON and OFF channels project to the excitatory neurons in layer 4 of the network. The strength of each synapse is determined by the neuron’s decoding weights, which in this model are defined by Gabor filters. These filters represent the classical receptive field of simple cells in Primary Visual Cortex (V1) and allow neurons to selectively respond to specific orientations and spatial frequencies [8, 9].

Each neuron is assigned a unique Gabor receptive field:

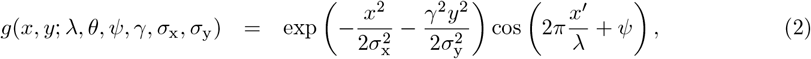

with rotated spatial coordinates:

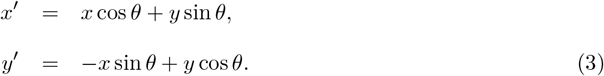

Each Gabor filter includes parameters for orientation (*θ*), phase (*ψ*), spatial frequency (*λ*), width (*σ*_x_ and *σ*_y_), and aspect ratio (*γ*). These are initialized to span a diverse and biologically realistic range across the population, as summarized in Table 3.

Both ON and OFF components of the Gabor filter contribute separately to the corresponding synaptic weights from ON and OFF LGN channels. All neurons use receptive fields the same size as the input patch (e.g., 16 × 16 pixels), simplifying implementation while preserving spatial selectivity.

Within the predictive coding framework, these Gabor filters determine the decoding weights that each neuron employs to reconstruct the stimulus. The total synaptic input is interpreted as the prediction error between the external input and the current estimate generated by the network. This interpretation follows the formulation introduced by Boerlin et al. [3], where the summed decoding weights of recently active neurons form the network’s estimate.

The selection of Gabor filter parameters in Table 3 is grounded in electrophysiological measurements from cat and macaque V1, where classical simple-cell receptive fields are well approximated by localized, oriented, and bandpass-tuned profiles [45, 46]. Orientation preferences were sampled uniformly to reflect the columnar diversity observed across the cortex. The chosen spatial frequency of 0.8 cycles per degree aligns with reported tuning peaks in primate V1 under natural viewing conditions [47,48]. Envelope widths and aspect ratios were set to capture the elongation and bandwidth properties typically observed in layer 4 simple cells. While Gabor filters remain a canonical choice for modelling early visual encoding, alternative frameworks have been proposed that also account for receptive field structure. These include log-polar mappings of visual space [49], sparse approximation methods inspired by V1 functional architecture [50], and invertible 2D log-Gabor wavelet bases that improve reconstruction fidelity and scale invariance [51]. These methods offer valuable perspectives on receptive field modelling, particularly in large-scale or human fMRI contexts. However, Gabor functions offer an analytically simple, biologically validated, and computationally tractable basis for initialising decoding weights in spiking models and are therefore well-suited for our simulation goals.

This pre-processing and decoding structure enables the network to transform incoming sensory signals into orientation- and phase-specific spike patterns, laying the groundwork for predictive representation in the subsequent dynamics. All pre-processing steps in the simulation were implemented using Python. The full pre-processing pipeline, along with simulation code, is publicly available at GitHub repository.

### 2.3 Network Dynamics

labelssec:network-dynamics The network is based on the predictive coding framework introduced by Boerlin et al. [3], where spikes are fired to reduce the mismatch between the sensory input and the network’s internal prediction. Each neuron’s membrane potential reflects the local prediction error. The network includes distinct excitatory and inhibitory populations with decoding weight matrices *G*_E_ and *G*_I_, respectively, and all connectivity respects Dale’s law.

To translate Denève’s spike-based architecture into a biologically grounded model, synaptic interactions are divided into two components: fast and slow. Fast connections are modeled as instantaneous synaptic inputs that update the membrane potential at each presynaptic spike, with current amplitudes scaled by the synaptic weights *W* . Slow connections, on the other hand, are implemented by low-pass filtering the spike trains, and the resulting current is injected into the membrane equation continuously at each simulation step. In the Denève model, synaptic interactions are assumed to be instantaneous, with synaptic time constants set equal to the simulation time step Δ *t*. In our implementation, we set the synaptic time constants to biologically realistic values, with excitatory synapses set to *τ*_E_ = 5 ms and inhibitory synapses set to *τ*_I_ = 10 ms. This adjustment does not significantly affect the network dynamics. Furthermore, refractory periods were incorporated enhance the biological grounding of the model and which limits excessive spiking, with *τ*_r,E_ and *τ*_r,I_ set to 2 ms and 0.5 ms, respectively.

The model in this study receives discrete spike trains from an LGN-like input population, denoted *S*_LGN_(*t*), providing a more physiologically grounded representation of sensory drive. Each neuron’s decoding weight (*G*_E_, *G*_I_) is defined by a biologically inspired Gabor filter, resulting in receptive fields that resemble those of layer 4 simple cells in V1. In contrast, Denève’s original formulation [3] assumes a continuous-valued input signal *c*(*t*), with synaptic weights typically initialized randomly or selected arbitrarily. The present model, therefore, more closely reflects cortical anatomy and feature selectivity.

#### Excitatory Dynamics

In Denève’s original predictive coding formulation, the membrane potential of excitatory neurons evolves according to:

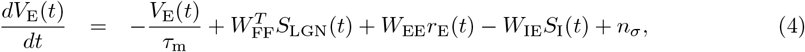

where *V*_E_(*t*) denotes the membrane potential of excitatory neurons, *τ*_m_ is the membrane time constant, and *n*_*σ*_ is a zero-mean Gaussian noise term. The weight matrices are defined as follows: *W*_FF_ = *G*_E_ for feedforward LGN input, and 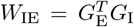 for inhibitory feedback. In addition, 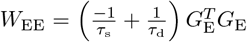 for recurrent excitatory feedback, where *τ*_d_ and *τ*_s_ are decoding and sensory time constants.

The firing rate *r*(*t*) is obtained by filtering the spike train *S*(*t*) with an exponential kernel:

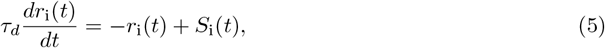

where *S*_i_(*t*) represents the spike train of neuron *i*. This filtered firing rate is then used to drive the slow synaptic feedback.

In the equivalent current-based model, the excitatory dynamics are implemented as:

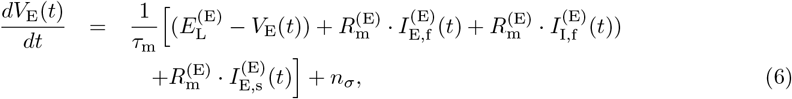

where 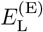 is the leak reversal potential, 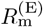 is the membrane resistance, and 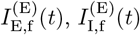, and 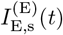 represent the fast excitatory, fast inhibitory, and slow excitatory synaptic input currents received by the excitatory population, respectively. The term *n*_*σ*_ adds zero-mean Gaussian noise to introduce variability in the membrane potential.

The evolution of the synaptic currents follows:

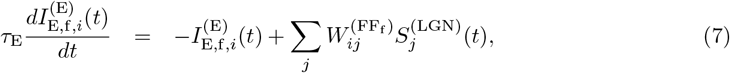

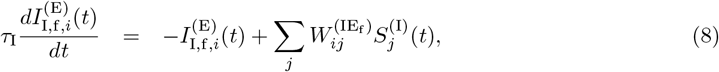

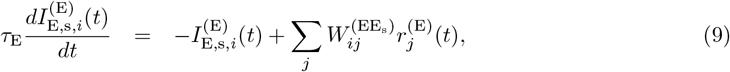

where *τ*_E_ and *τ*_I_ are the excitatory and inhibitory synaptic time constants, respectively. The synaptic weight matrices 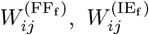, and 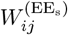 define the connections from LGN, inhibitory, and excitatory populations, respectively. 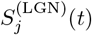 and 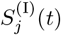 represent spike trains from the LGN and inhibitory neurons, while 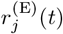 denotes the instantaneous firing rate of excitatory neuron *j*. All currents are defined for postsynaptic excitatory neuron *i*. The synaptic weight matrices are adapted from Deneve’s original formulation, initialized here using Gabor decoding weights.

#### Inhibitory Dynamics

The inhibitory membrane potential in Denève’s framework evolves as:

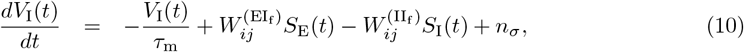

where *V*_I_(*t*) is the membrane potential of inhibitory neurons, *τ*_m_ is the membrane time constant, and *n*_*σ*_ is a zero-mean Gaussian noise term. 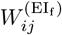 and 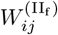 are the synaptic weights from excitatory and inhibitory neurons, respectively. The spike trains *S*_E_(*t*) and *S*_I_(*t*) represent the input from presynaptic excitatory and inhibitory populations.

In the equivalent current-based model, the inhibitory dynamics are described by:

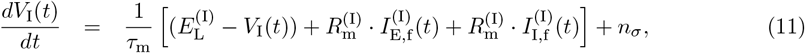

where 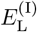 is the leak potential, 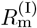 is the membrane resistance, and 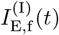 and 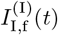 are the fast excitatory and inhibitory synaptic input currents to the inhibitory population, respectively.

The synaptic currents evolve according to:

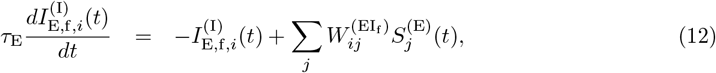

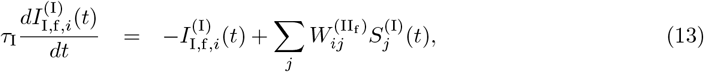

where 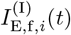 and 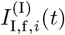 are the fast synaptic currents received by inhibitory neuron *i* from excitatory and inhibitory populations, respectively.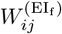 and 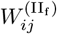 denote the synaptic weight matrices, while 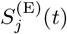 and 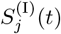 represent the spike train of excitatory and inhibitory neuron neuron *j*, respectively.

This formulation allows inhibitory neurons to dynamically regulate the excitatory population by responding rapidly to incoming spike activity, following Denève’s predictive coding framework.

### 2.4 Biophysical Rescaling and Threshold Matching

To express voltage and synaptic activity in biological units, each neuron’s internal membrane potential *V*_i_(*t*) is linearly rescaled:

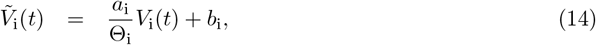

where *a*_i_ controls the amplitude (e.g., 25 mV) and *b*_i_ sets the resting potential (e.g., −80 mV). The biologically scaled spiking threshold becomes:

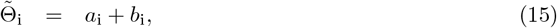

and the synaptic weights are rescaled as:

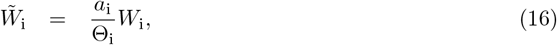

allowing membrane potentials to evolve in millivolts and produce realistic thresholds (e.g., −55 mV), while ensuring that synaptic weights yield typical cortical postsynaptic potential (PSP) amplitudes.

To preserve balance as the network size *N* increases, the following scaling rules are applied:

- 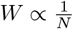 (normalizes total input),
- 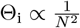 (ensures sufficient drive for spiking).

These rules maintain stable membrane fluctuations, realistic firing thresholds, and consistent input strength in large populations. Feedforward weights scale with the norm of the decoding kernels, while recurrent weights remain invariant to their amplitude.

By aligning model parameters with biologically observed values, such as resting potential, threshold, and synaptic strength, this transformation ensures that membrane voltages and synaptic inputs fall within physiologically realistic ranges.

Taken together, the excitatory and inhibitory dynamics described above enable the network to sustain biologically realistic activity while performing continuous real-time prediction error correction [3]. To achieve this, all model parameters were aligned with biologically observed values such as resting potential, threshold, and synaptic strength, ensuring that membrane voltages and synaptic inputs fall within physiologically realistic ranges.

The synaptic weights were derived from decoding weights *G*_E_ and *G*_I_, instantiated as Gabor filters to reflect the orientation and phase tuning of layer 4 simple cells. Membrane and leak parameters, refractory periods, and noise levels were set to yield realistic firing rates and membrane fluctuations, consistent with values observed in cortical circuits [10, 11, 26].

### 2.5 Functional Measures and Network Readouts

To evaluate the network’s functionality, we employed several quantitative measures that capture selectivity, irregularity, synchrony, and balance. These were computed from spike trains, filtered synaptic currents, and response statistics under different stimulus conditions.

#### Orientation and Phase Bias Indices (*π*_*θ*_ and *π*_*ψ*_)

Orientation selectivity was quantified using the Orientation Bias Index (OBI),

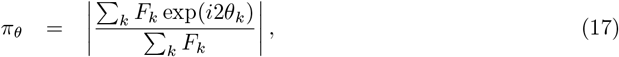

where *F*_*k*_ denotes the neural response to a stimulus with orientation *θ*_*k*_. Similarly, phase selectivity was measured with the Phase Bias Index (PBI),

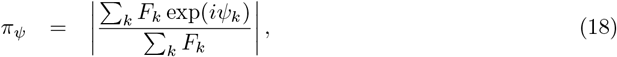

where *ψ*_*k*_ represents the phase of each stimulus. These indices range from 0 to 1, with higher values indicating stronger tuning [52, 53].

#### Modulation Ratio (*M*_R_)

To assess phase sensitivity, we computed the *M*_R_ ratio:

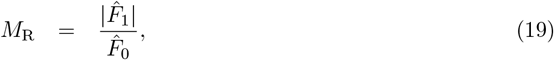

where 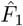 is the amplitude of the first harmonic of the phase tuning curve and 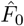 is the mean response. Neurons with *M*_R_ *>* 1 are classified as simple cells; those with *M*_R_ *<* 1 are complex cells [54–56].

#### Spike Train Irregularity 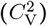

Spike timing irregularity was measured using the squared coefficient of variation of Inter-Spike Interval (ISI) (*δ*):

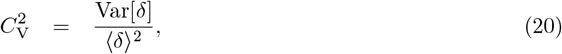

where ⟨·⟩ is the mean, and Var denotes variance. Values 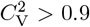 indicate irregular (Poisson-like) firing [57].

#### Synchrony 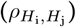

Pairwise synchrony was measured using the Pearson correlation of binned spike trains:

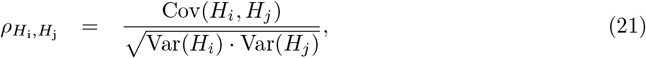

where *H*_*i*_ and *H*_*j*_ are spike count time series for neurons *i* and *j* (binned at 10 ms). Higher values reflect greater spike-time coordination [26].

#### Excitatory-Inhibitory (E-I) Balance

We used three current-based measures to evaluate E-I balance from smoothed (10 ms) current traces:

##### 1 – Slope (*m*)

The linear scaling between inhibitory and excitatory currents:

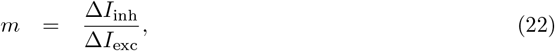

A slope near 1 suggests proportional E–I coupling.

##### 2 – Correlation (*ρ*)

The Pearson correlation coefficient between excitatory and inhibitory current traces:

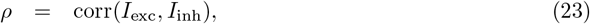

Higher *ρ* values reflect tighter temporal coordination.

##### 3 – Balance Index (*β*)

Following the approach of Ahmadian and Miller [58], we computed a normalized balance index to quantify the net input deviation from perfect balance. The mean excitatory input is:

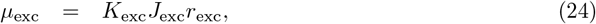

where *K*_exc_ is the number of inputs, *J*_exc_ the mean synaptic weight, and *r*_exc_ the presynaptic firing rate. The total mean input current is given by:

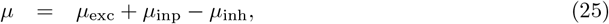

and the balance index is defined as:

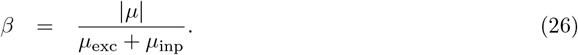

This ratio quantifies the deviation of total input from perfect cancellation. Values *β* ≪ 1 indicate tight balance, whereas *β >* 0.1 signal imbalance. This metric was computed per neuron based on the recorded synaptic current traces.

## 3 Results

In this section, a detailed characterization of network structure, activity, and input-output transformations across different stimulus conditions are presented. The architectural organization of the model is first described, which reflects key anatomical and functional features of the V1, including orientation-and phase-specific feedforward connectivity and structured recurrent interactions. We then verify that the Poisson-driven LGN input exhibits expected statistical properties when receiving noise and grating input stimuli. Following this, the cortical network response is examined, highlighting spiking patterns, membrane potential distributions, and input current statistics for excitatory and inhibitory populations. Particular emphasis is placed on the emergence of excitatory-inhibitory balance under random and structured inputs, both at the level of single neurons and across the population. We further explore how the network dynamics adapt to varying contrast levels and systematically quantify orientation and phase selectivity. Finally, we assess the robustness of the network’s ability to reconstruct input stimuli under increasing noise levels, providing insights into the fidelity of sensory representations.

### 3.1 Network Structure

The model network was designed to reflect key anatomical and functional properties of V1, with neurons organized by their preferred orientation and phase, and connected via structured feedforward and recurrent connectivity.

Fig. 3A visualizes the feedforward LGN-to-cortex connections, modeled as Gabor filters. Each filter reflects the receptive field of a cortical neuron, parameterized by its preferred spatial frequency, orientation, and phase. This configuration supports orientation- and phase-specific selectivity.

**Fig 3.**
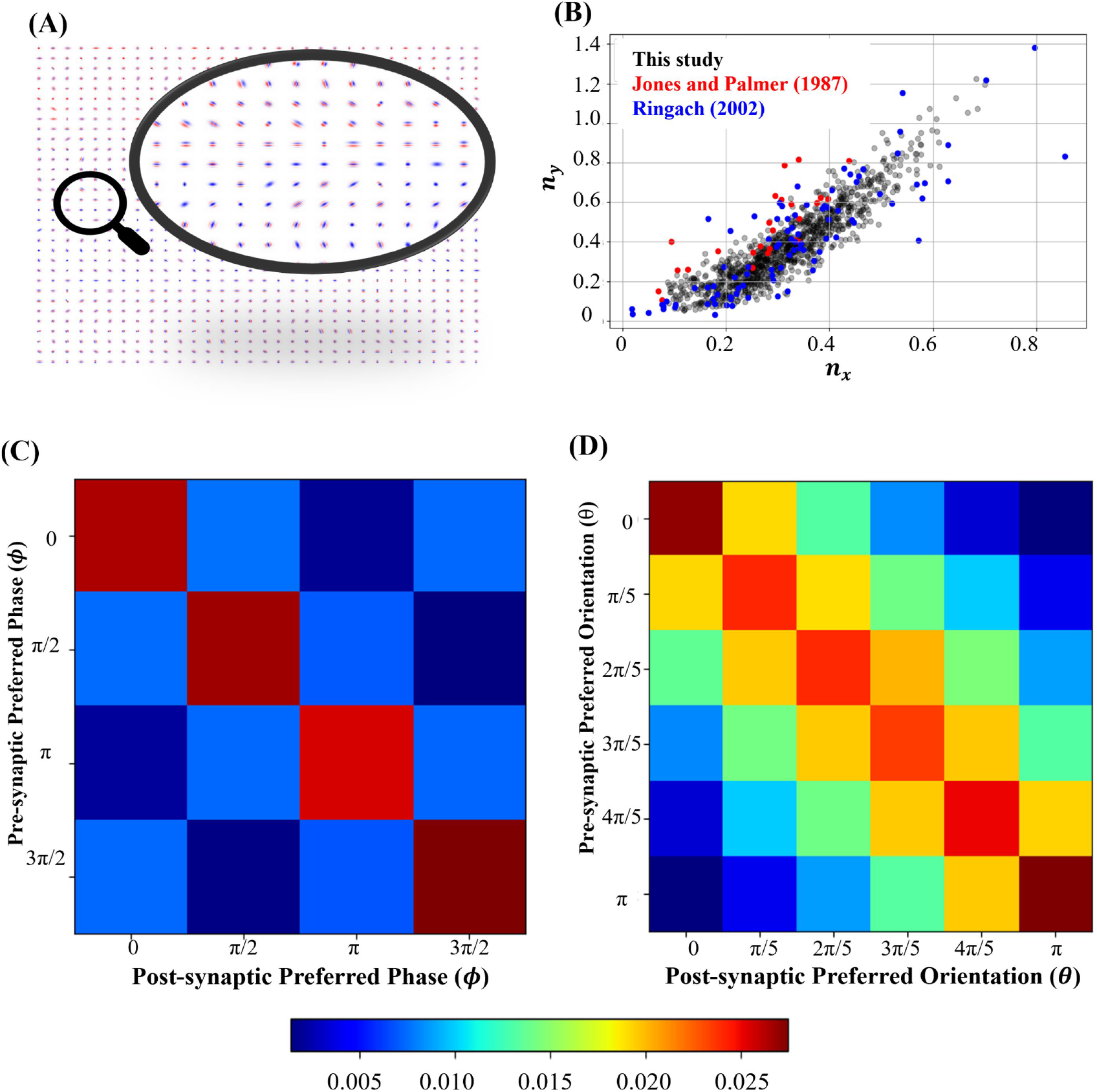
Structure and feature-dependent connectivity of the modeled cortical network. **(A)** Feedforward receptive fields from LGN to cortical neurons are modeled as Gabor filters, with each vector field indicating local orientation and phase preference. The inset highlights the diversity and spatial distribution of these filters. **(B)** Comparison of receptive field spatial frequency structure from this study (black) with biological data from Jones and Palmer [8] (red) and Ringach [9] (blue). Each point represents a neuron’s Gabor envelope, plotted in terms of *n*_*x*_ and *n*_*y*_. **(C)** Phase-dependent connectivity matrix showing average synaptic strength between neuron pairs as a function of their preferred phase. Strongest connectivity occurs when pre- and post-synaptic phases are aligned. **(D)** Orientation-dependent connectivity matrix showing synaptic strength between neurons as a function of orientation difference. Neurons with similar orientation preferences form stronger connections, indicating feature-specific lateral structure.

To quantitatively evaluate the structure of receptive fields across the population, the shape of each fitted Gabor envelope was analyzed using the parameters *n*_*x*_ = *σ*_x_*f* and *n*_*y*_ = *σ*_y_*f*, where *σ*_x_ and *σ*_y_ are the standard deviations of the Gaussian envelope along the horizontal and vertical axes, and *f* is the spatial frequency. As shown in Fig. 3B, receptive fields from our model (black) fall within the range of those measured experimentally in cat V1 by Jones and Palmer [8] (red) and Ringach [9] (blue). The strong positive correlation between *n*_*x*_ and *n*_*y*_ is consistent with known biological constraints on aspect ratio and spatial structure.

The network also incorporates structured recurrent connectivity. Fig. 3C shows the average synaptic weight between neurons with identical orientation preference but differing in phase. Stronger connectivity is observed along the diagonal, indicating phase-specific excitation. Also, Fig. 3D presents connectivity among neurons sharing the same phase but differing in orientation. Connections are strongest between similarly tuned neurons and decay with increasing orientation difference, reflecting a known anatomical motif that supports orientation-specific integration in cortical circuits.

### 3.2 Network Response to Noise Input

To establish a realistic baseline regime, we first simulated cortical responses to spatially unstructured noise input from the LGN. This input was generated by presenting a randomized stimulus image (see inset in Fig. 4A), where ON-center and OFF-center LGN neurons (*N*_LGN,ON_ = 256 and *N*_LGN,OFF_ = 256) responded via a Poisson process. Spike rasters revealed stochastic but sustained firing across both populations (Fig. 4A), with ON cells firing at higher average rates due to stimulus bias.

**Fig 4.**
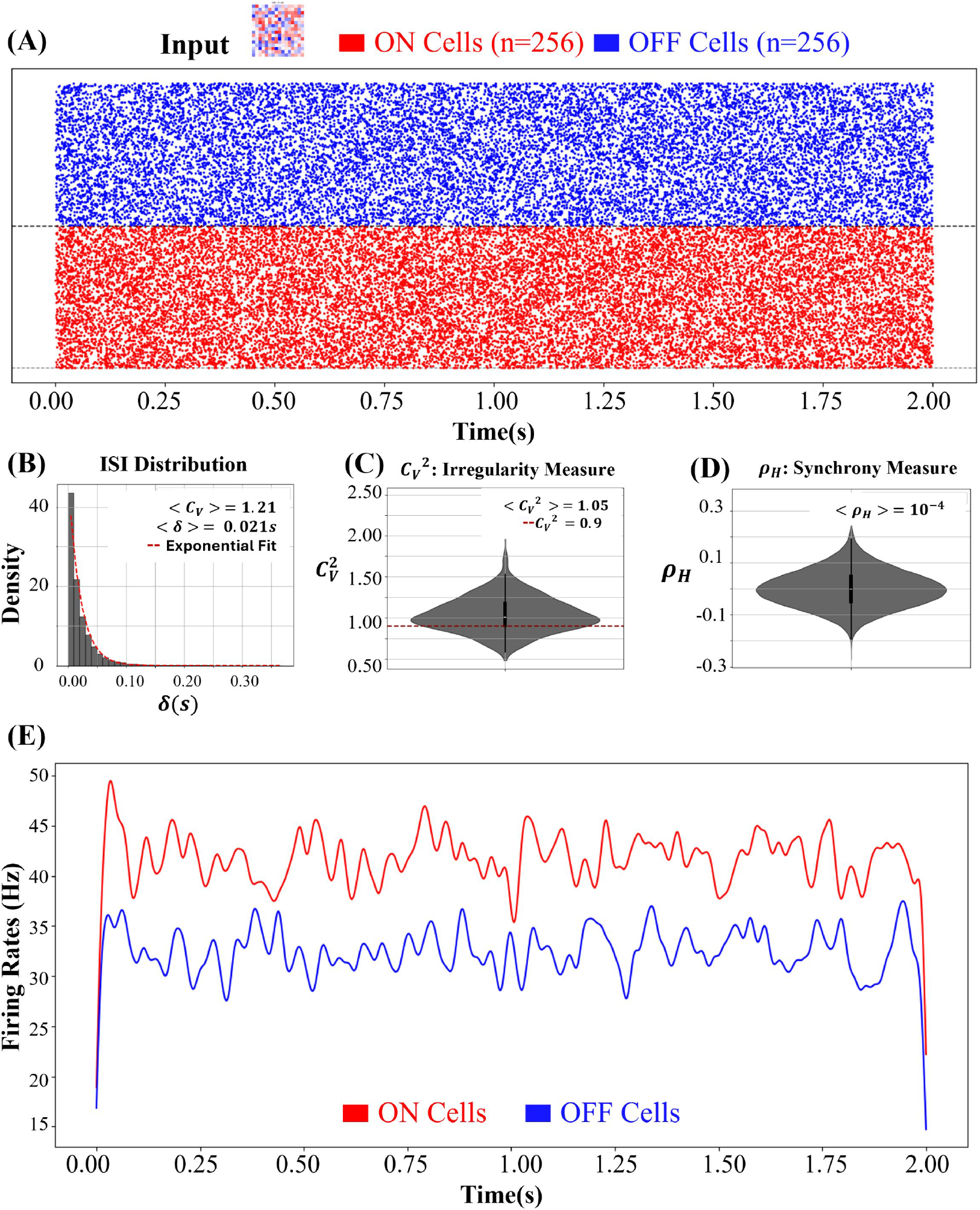
Poisson LGN input under noise stimulus and population-level statistics. **(A)** Raster plot of ON (red) and OFF (blue) LGN neurons (*N*_LGN,ON_ = 256 and *N*_LGN,OFF_ = 256). Inset: stimulus image. **(B)** ISI distribution with exponential fit; population mean: ⟨*C*_V_⟩ = 1.21. **(C)** Irregularity index 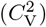, with 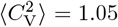 . Dashed line marks irregular threshold 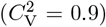 . **(D)** Spike histogram correlation 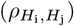 centered near zero, 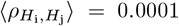, indicating low synchrony. **(E)** Firing rate profiles for ON and OFF populations, showing higher activity in ON neurons.

Across the LGN population, ISI distributions closely followed an exponential profile, with a population-wide mean coefficient of variation ⟨*C*_V_⟩ = 1.21 (Fig. 4B). The irregularity index 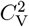, which quantifies local spike train variability, showed a population mean of 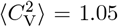, exceeding the irregularity threshold of 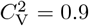 (Fig. 4C). Spike histogram correlations 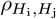 were centered near zero, with 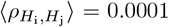 (Fig. 4D), confirming low synchrony across neurons. Firing rate profiles (Fig. 4E) showed sustained activity over time, with the ON population exhibiting consistently higher rates than the OFF population.

The downstream response in cortical layer 4 is shown in Fig. 5. Raster plots (Fig. 5A) demonstrate sparse, irregular spiking across the excitatory (red) and inhibitory (blue) populations, filtered to include only neurons with more than 10 spikes. Violin plots of membrane potential distributions (Fig. 5B) show overlapping subthreshold dynamics. Synaptic current distributions (Fig. 5C) indicate balanced excitatory and inhibitory input strengths within each population.

**Fig 5.**
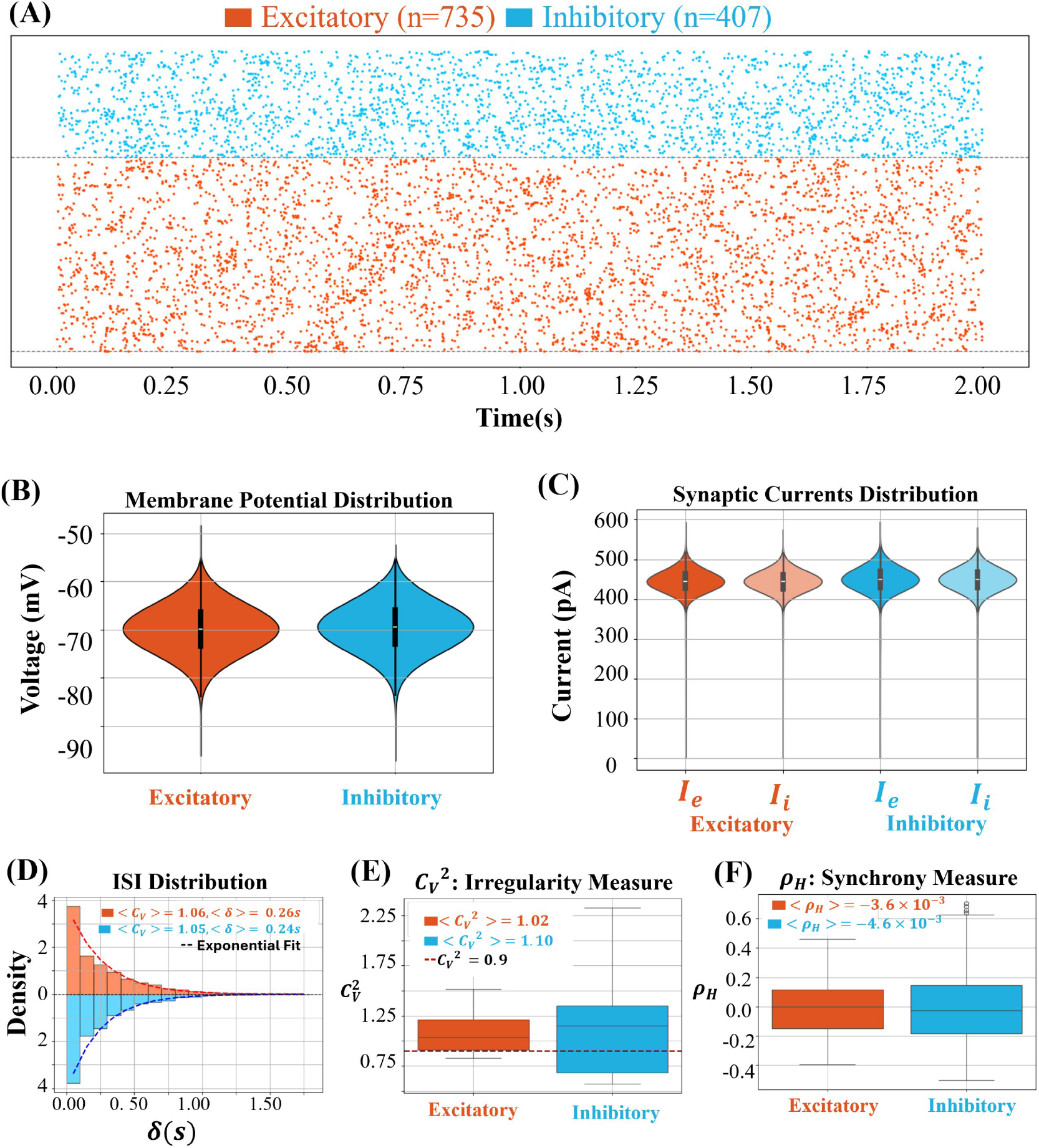
Cortical population response to noise input. **(A)** Raster plot of excitatory (red) and inhibitory (blue) neurons, filtered for *>* 10 spikes. **(B)** Membrane potential (*V*_m_(*t*)) distributions showing subthreshold overlap. **(C)** Synaptic current distributions for excitatory (*I*_E_(*t*)) and inhibitory (*I*_I_(*t*)) input, grouped by population. **(D)** ISI distributions with population means: ⟨*C*_V_⟩ = 1.06 (exc), ⟨*C*_V_⟩ = 1.05 (inh). **(E)** Irregularity index 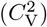 distributions with 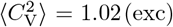, 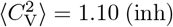 ; dashed line marks 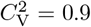 . **(F)** Synchrony measure 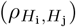 distributions with 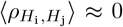, indicating low synchrony in spiking activity. Violin plots show full data distribution; overlaid box plots indicate inter-quartile range and median, with horizontal lines marking mean values.

Both populations exhibited near-Poisson spike timing, with population means ⟨*C*_V_⟩ = 1.06 for excitatory neurons and ⟨*C*_V_⟩ = 1.05 for inhibitory neurons, as shown in the ISI histograms (Fig. 5D). Irregularity indices 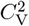 exceeded the threshold of 0.9 in both groups (Fig. 5E), confirming irregular spiking under baseline noise, with inhibitory neurons showing slightly greater variability 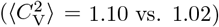. Spike histogram correlations 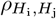 remained close to zero (Fig. 5F), consistent with weak synchrony across population activity in response to unstructured input.

To examine excitatory-inhibitory balance at the single-cell level, we analyzed representative neurons from both populations (Fig. 6). Scatter plots of excitatory versus inhibitory input currents (Fig. 6A,B) revealed strong negative correlations in both the excitatory and inhibitory neural populations. For the excitatory neurons, the linear fit had a slope of *m* = 0.73, a Pearson correlation coefficient of *ρ* = 0.85, and a balance index of *β* = 0.01. For the inhibitory neurons, the slope was *m* = 0.80, with *ρ* = 0.94 and *β* = 0.01. These results confirm dynamically matched excitation and inhibition at the single-neuron level, supporting a tightly balanced E-I regime under unstructured noisy input. The membrane potential traces (Fig. 6C,D) show subthreshold fluctuations interspersed with occasional spikes. The input current traces (Fig. 6E,F) further illustrate opposing fluctuations in *I*_E_(*t*) and *I*_I_(*t*), with relatively stable net current (*I*_n_(*t*)), reflecting precise dynamic cancellation. Variability measures indicate a slightly larger standard deviation in excitation than inhibition, consistent across cell types.

**Fig 6.**
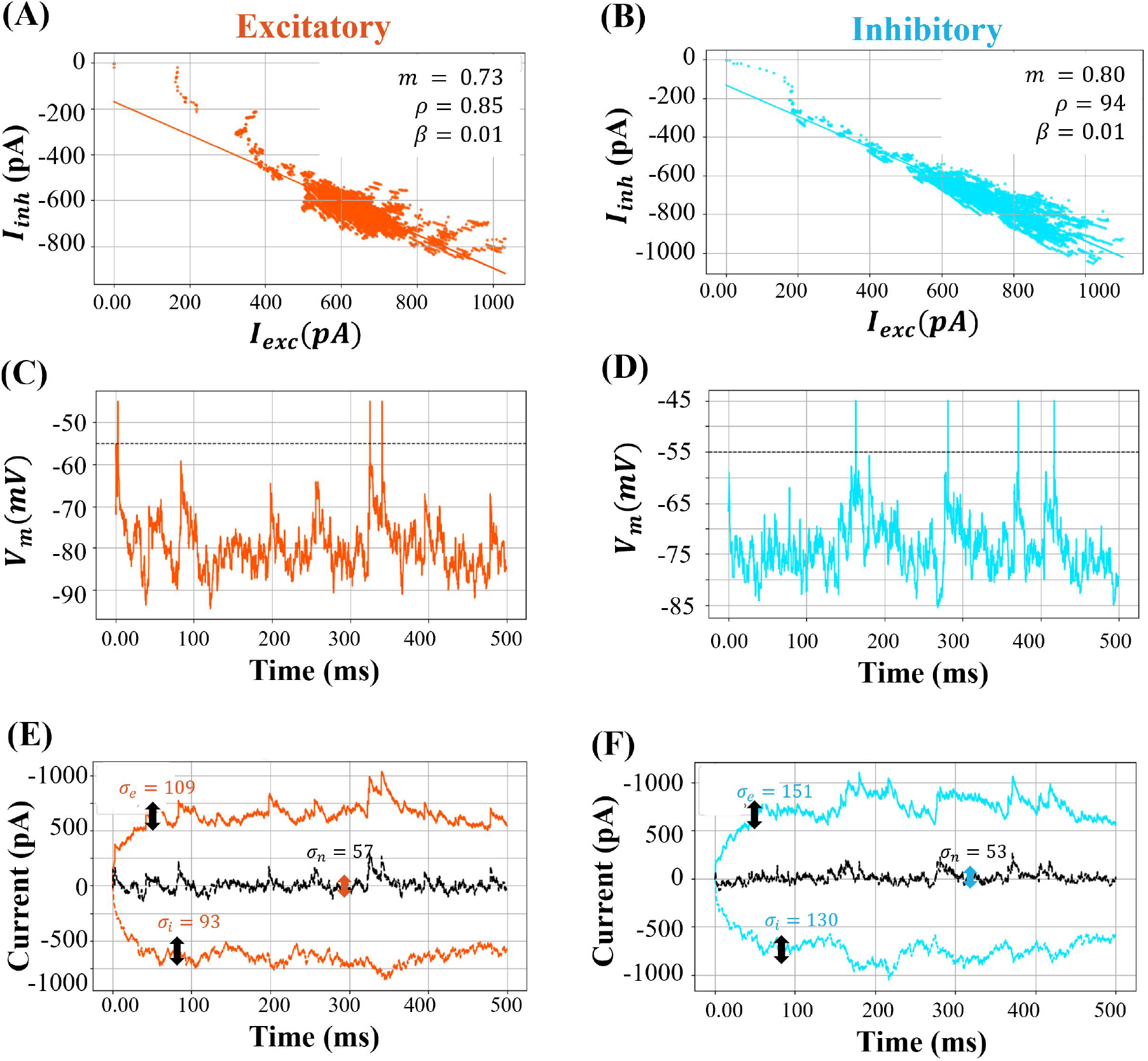
Single-neuron E-I balance under noise input. **(A, B)** *I*_E_(*t*) vs. *I*_I_(*t*) scatter plots for representative excitatory and inhibitory neurons. Linear fits show strong anti-correlation (*m* = 0.73, *ρ* = 0.85 for excitatory; *m* = 0.80, *ρ* = 0.94 for inhibitory), with low balance index (*β* = 0.01). **(C, D)** Membrane potential (*V*_m_(*t*)) traces for each neuron with spike threshold (dashed line). **(E, F)** Time courses of excitatory (*I*_E_(*t*)), inhibitory (*I*_I_(*t*)), and net current (*I*_n_(*t*) = *I*_E_(*t*) + *I*_I_(*t*)). Variability is quantified by standard deviations: *σ*_*e*_ = 109 pA, *σ*_*i*_ = 93 pA, *σ*_*n*_ = 57 pA for excitatory; *σ*_*e*_ = 151 pA, *σ*_*i*_ = 130 pA, *σ*_*n*_ = 53 pA for inhibitory.

### 3.3 Network Response to Grating Input

Cortical responses to structured input were examined by presenting a grating input to the LGN. Raster plots of ON-center and OFF-center LGN neurons (*N*_LGN,ON_ = 256 and *N*_LGN,OFF_ = 256) in Fig. 7A show stimulus-locked synchrony within subpopulations tuned to the grating features. The structured input resulted in visible banded patterns of spike activity, corresponding to the spatial modulation of the grating.

**Fig 7.**
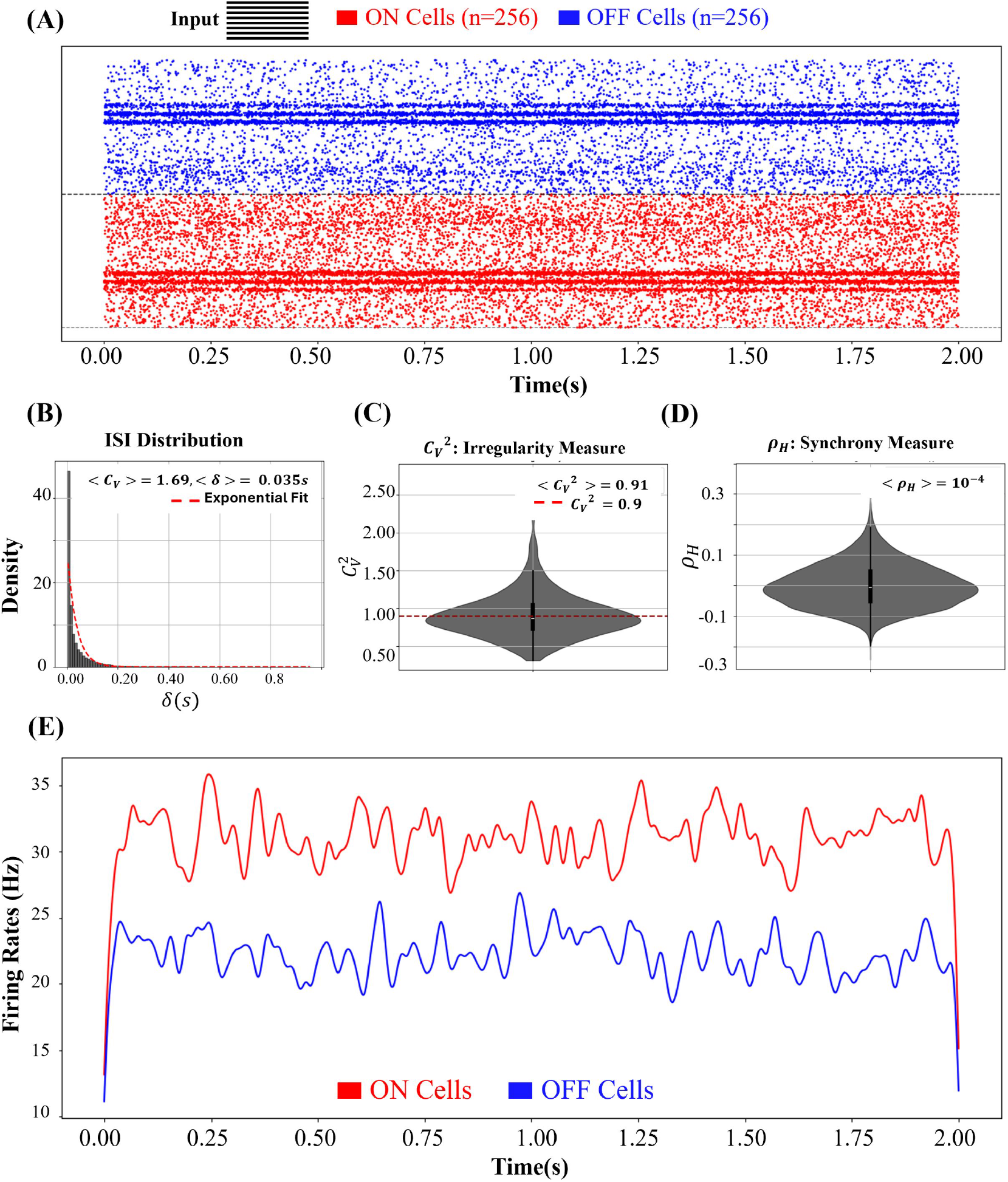
Poisson LGN input and statistical characterization for grating stimulus. **(A)** Raster plot of ON (red) and OFF (blue) LGN neurons (*N*_LGN,ON_ = 256 and *N*_LGN,OFF_ = 256) in response to grating input. Inset: stimulus image. **(B)** ISI distribution with exponential fit; population mean: ⟨*C*_V_⟩ = 1.69. **(C)** Irregularity index 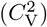 with 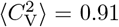 . Dashed line marks the irregularity threshold at 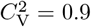 . **(D)** Spike histogram correlation 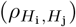 centered near zero, 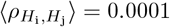, suggesting weak synchrony. **(E)** Firing rate profiles for ON and OFF populations during grating input, showing higher activation in ON neurons.

Across the LGN population, ISI distributions exhibited a sharper decay compared to the noise input, with a higher population-wide coefficient of variation ⟨*C*_V_⟩ = 1.69 (Fig. 7B). The irregularity index 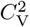 averaged 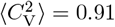 (Fig. 7C), close to the irregularity threshold of 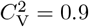, suggesting a borderline transition between irregular and more regular spiking dynamics. Spike histogram correlations 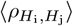 remained centered around zero (Fig. 7D), indicating low pairwise synchrony despite the presence of spatial structure in the stimulus. The firing rate profiles (Fig. 7E) showed that ON neurons maintained consistently higher firing rates than OFF neurons, reflecting their preferential activation by the grating.

The downstream cortical response to the grating input is shown in Fig. 8. Raster plots (Fig. 8A) depict spiking activity for excitatory (*N*_E_ = 615) and inhibitory (*N*_I_ = 235) neurons, filtered to include cells with more than 10 spikes. Both populations displayed structured, periodic firing patterns reflecting the spatially modulated input.

**Fig 8.**
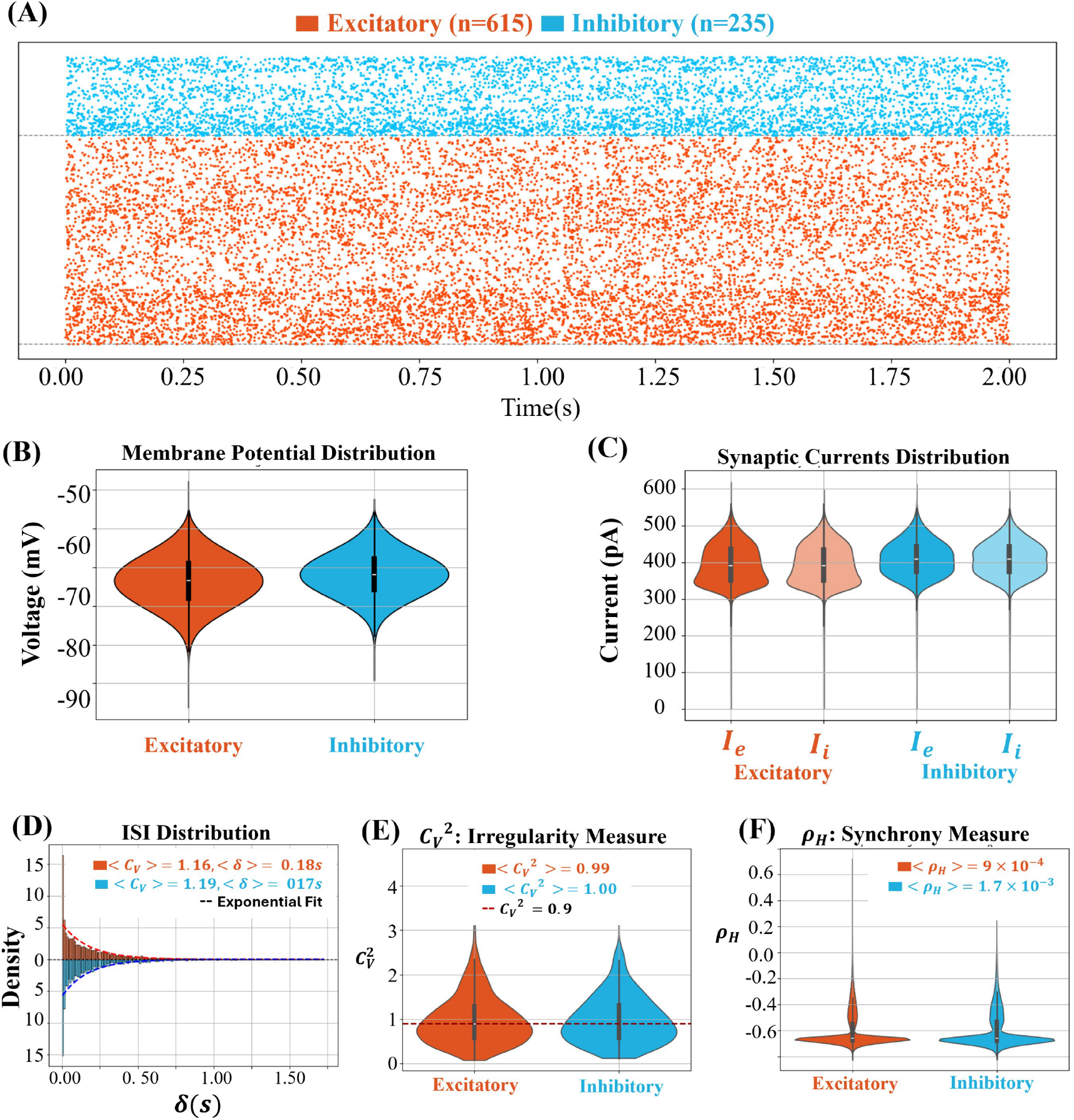
Cortical population response to grating input. **(A)** Raster plot of excitatory (red) and inhibitory (blue) neurons during grating stimulation, filtered for neurons with *>* 10 spikes. **(B)** Membrane potential (*V*_m_(*t*)) distributions showing similar subthreshold profiles across populations. **(C)** Synaptic current distributions for excitatory (*I*_E_(*t*)) and inhibitory (*I*_I_(*t*)) inputs, grouped by population. **(D)** ISI distributions with population means: ⟨*C*_V_⟩ = 1.16, ⟨*δ*⟩ = 0.18 s (exc); ⟨*C*_V_⟩ = 1.19, ⟨*δ*⟩ = 0.17 s (inh). **(E)** Irregularity index distributions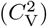, with population means: 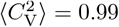 (exc), 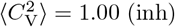 dashed line marks 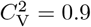 . **(F)** Synchrony measure 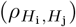 based on histogram correlations: 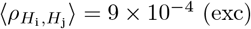,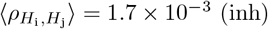.

Violin plots of membrane potential distributions (Fig. 8B) showed overlapping subthreshold activity between excitatory and inhibitory populations. Synaptic current distributions (Fig. 8C) showed balanced excitatory and inhibitory input within each population, with slightly broader distributions for excitatory inputs. The ISI distributions (Fig. 8D) exhibited moderate variability, with population means of ⟨*C*_V_⟩ = 1.16 and ⟨*δ*⟩ = 0.18 s for excitatory neurons, and ⟨*C*_V_⟩ = 1.19, ⟨*δ*⟩ = 0.17 s for inhibitory neurons.

Irregularity indices 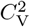 (Fig. 8E) were close to unity in both populations, with population means 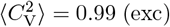 and 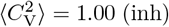, indicating near-Poisson spike timing. Synchrony measures 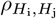 (Fig. 8F) remained low across populations, with 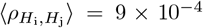 for excitatory and 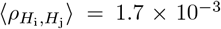 for inhibitory neurons, confirming weak synchrony across neurons under grating input.

Single-neuron analyses further confirmed the emergence of tight excitatory-inhibitory balance under structured input (Fig. 9). For a representative excitatory neuron, excitatory and inhibitory input currents were strongly anti-correlated, with a regression slope *m* = 0.92, Pearson correlation coefficient *ρ* = 0.96, and balance index *β* = 0.01 (Fig. 9A). For the inhibitory neuron, the corresponding values were *m* = 0.88, *ρ* = 0.97, and *β* = 0.02 (Fig. 9B).

**Fig 9.**
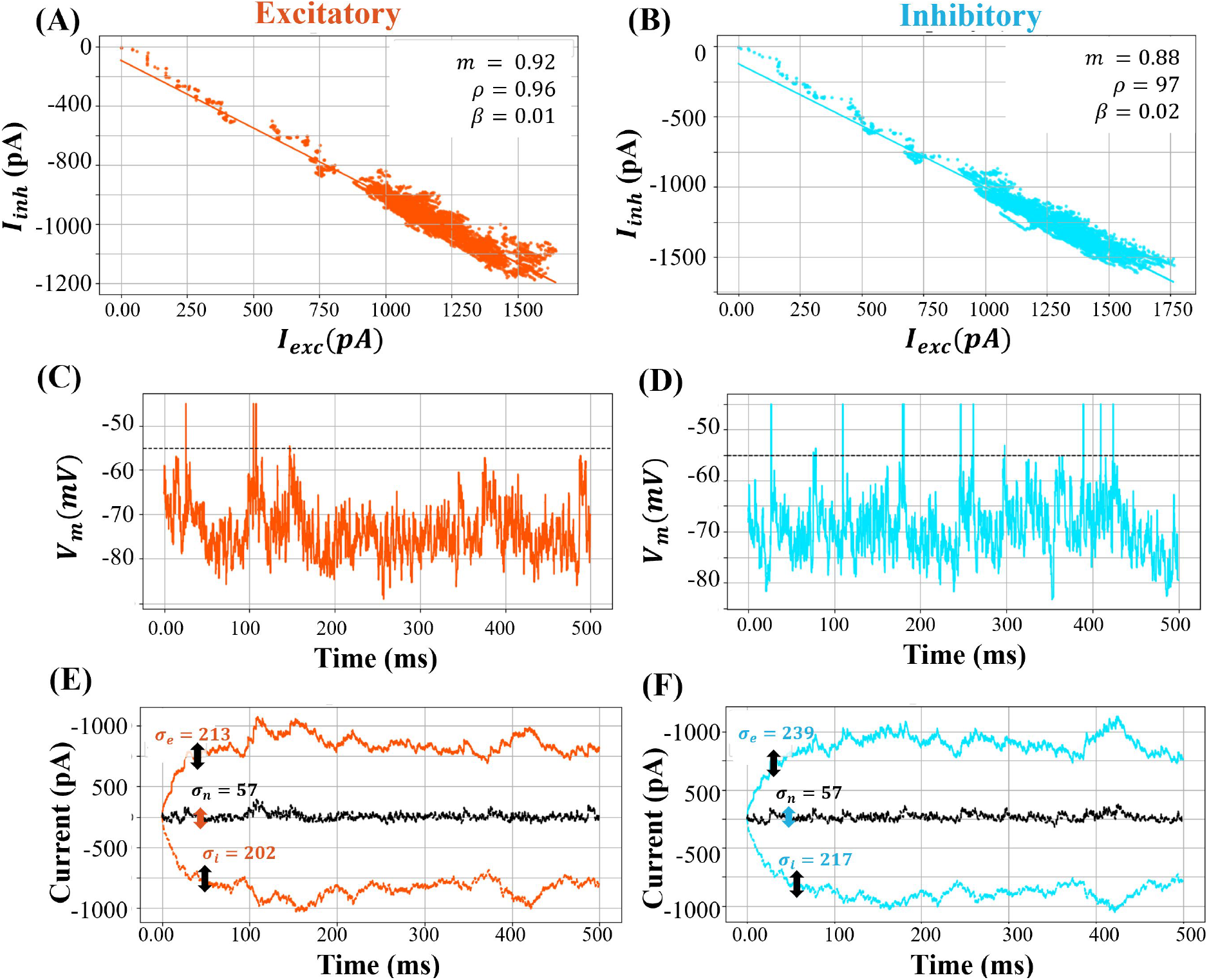
Single-neuron E-I balance under grating input. **(A, B)** *I*_E_(*t*) vs. *I*_I_(*t*) scatter plots for representative excitatory and inhibitory neurons. Linear fits show strong anti-correlation (*m* = 0.92, *ρ* = 0.96 for excitatory; *m* = 0.88, *ρ* = 0.97 for inhibitory), with low balance indices (*β* = 0.01, 0.02). **(C, D)** Membrane potential (*V*_m_(*t*)) traces for each neuron with spike threshold (dashed line). **(E, F)** Time courses of excitatory (*I*_E_(*t*)), inhibitory (*I*_I_(*t*)), and net current (*I*_n_(*t*) = *I*_E_(*t*)+*I*_I_(*t*)). Variability is quantified by standard deviations: *σ*_*e*_ = 213 pA, *σ*_*i*_ = 202 pA, *σ*_*n*_ = 57 pA for excitatory; *σ*_*e*_ = 239 pA, *σ*_*i*_ = 217 pA, *σ*_*n*_ = 57 pA for inhibitory.

Subthreshold membrane potential traces (Fig. 9C, D) showed regular depolarization events synchronized with the grating input. Input current traces (Fig. 9E, F) exhibited tight temporal alignment between *I*_E_(*t*) and *I*_I_(*t*), with small net currents *I*_n_(*t*) = *I*_E_(*t*) + *I*_I_(*t*) and low variability. For the excitatory neuron, current standard deviations were *σ*_*e*_ = 213 pA, *σ*_*i*_ = 202 pA, and *σ*_*n*_ = 57 pA; for the inhibitory neuron, they were *σ*_*e*_ = 239 pA, *σ*_*i*_ = 217 pA, and *σ*_*n*_ = 57 pA.

### 3.4 Comparison of Excitatory-Inhibitory Balance under Noise and Grating Inputs

To assess the robustness of E-I balance across different input regimes, we compared network dynamics under random noise versus structured grating stimulation. Fig. 10 summarizes key population-level balance measures for both conditions.

**Fig 10.**
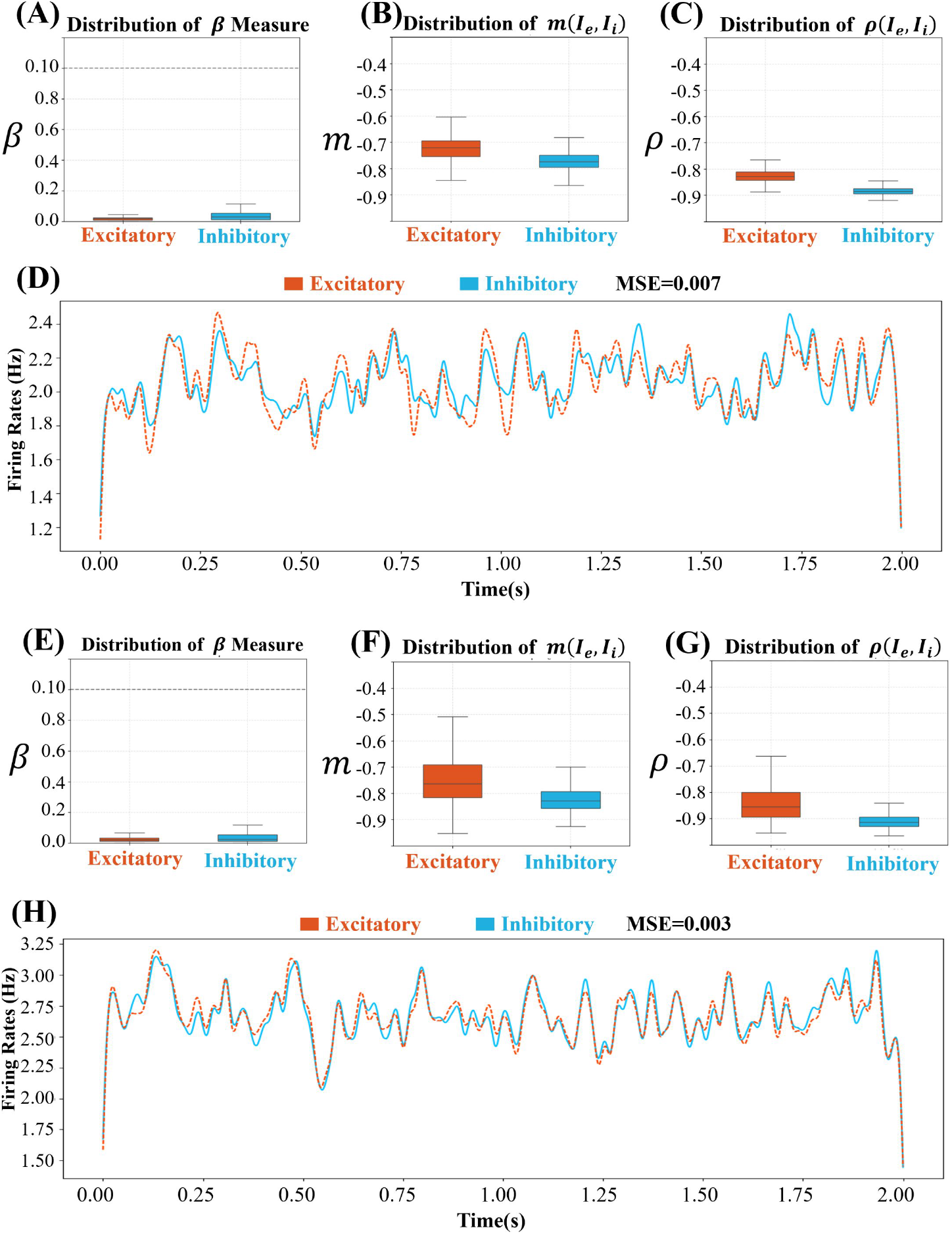
E-I balance under noise versus grating input (population-level comparison). Excitatory neurons are shown in red, and inhibitory neurons in blue throughout all panels. **(A–C)** Summary statistics under noise input: **(A)** Distribution of the balance index *β* across neurons. The dashed line at *β* = 0.1 marks the threshold separating tightly balanced (*β <* 0.1) from loosely balanced (*β >* 0.1) neurons. **(B)** Distribution of slopes *m*(*I*_E_(*t*), *I*_I_(*t*)) from linear fits between excitatory and inhibitory currents. **(C)** Distribution of Pearson correlation coefficients *ρ*(*I*_E_(*t*), *I*_I_(*t*)). **(D)** Firing rate alignment under noise input: excitatory population (red, dashed), scaled by *a* = 4 (reflecting the population size ratio), closely matches inhibitory population (blue), with MSE = 0.007. **(E–G)** Corresponding statistics under grating input: **(E)** Distribution of *β*, showing a wider spread. **(F)** Distribution of slopes *m*(*I*_E_(*t*), *I*_I_(*t*)). **(G)** Distribution of Pearson correlation coefficients *ρ*(*I*_E_(*t*), *I*_I_(*t*)). **(H)** Firing rate alignment under grating input: scaled excitatory and inhibitory rates remain tightly matched, with MSE = .003.

Panels (A–D) show balance statistics for the noise-driven regime. The distribution of the balance index *β* across neurons (Fig. 10A) reveals tight E-I balance throughout the population, with values consistently below the *β* = 0.1 threshold that separates tightly balanced (*β* ≪ 1) from loosely balanced neurons. The distribution of regression slopes *m*(*I*_E_(*t*), *I*_I_(*t*)) from linear fits between excitatory and inhibitory input currents (Fig. 10B) centers near −1, indicating that excitation and inhibition are dynamically matched in both magnitude and direction. Similarly, the distribution of Pearson correlation coefficients *ρ*(*I*_E_(*t*), *I*_I_(*t*)) (Fig. 10C) shows strong negative co-fluctuations across the population.

Population-averaged firing rates under noise input (Fig. 10D) further confirm E-I alignment: the excitatory firing rate (red dashed trace), scaled by a factor *a* = 4 (reflecting the population size ratio), closely matches the inhibitory rate (blue trace), with a low mean squared error (MSE = 0.007).

Panels (E–H) illustrate the same balance measures for the grating-driven regime. The distribution of *β* values (Fig. 10E) exhibits a wider spread and slightly higher mean, indicating that balance is preserved under structured input. The slopes *m*(*I*_E_(*t*), *I*_I_(*t*)) (Fig. 10F) remain consistently negative, and the correlation coefficients *ρ*(*I*_E_(*t*), *I*_I_(*t*)) (Fig. 10G) reflect continued tight co-fluctuations between excitatory and inhibitory input.

Firing rate dynamics under grating input (Fig. 10H) again demonstrate close tracking between scaled excitatory and inhibitory populations, with a similarly low MSE of 0.003. These results confirm that the cortical network maintains robust E-I balance under random and structured inputs.

### 3.5 Contrast-Dependent Excitatory-Inhibitory Dynamics

To investigate how stimulus contrast modulates cortical dynamics, we presented a temporally modulated Gabor stimulus at three contrast levels: low (0.25), medium (0.75), and high (1.25). The stimulus orientation was fixed while contrast changed every 500 ms, as indicated by grayscale shading in Fig. 11A. Only neurons that fired more than 10 spikes during the trial were included in the analysis.

**Fig 11.**
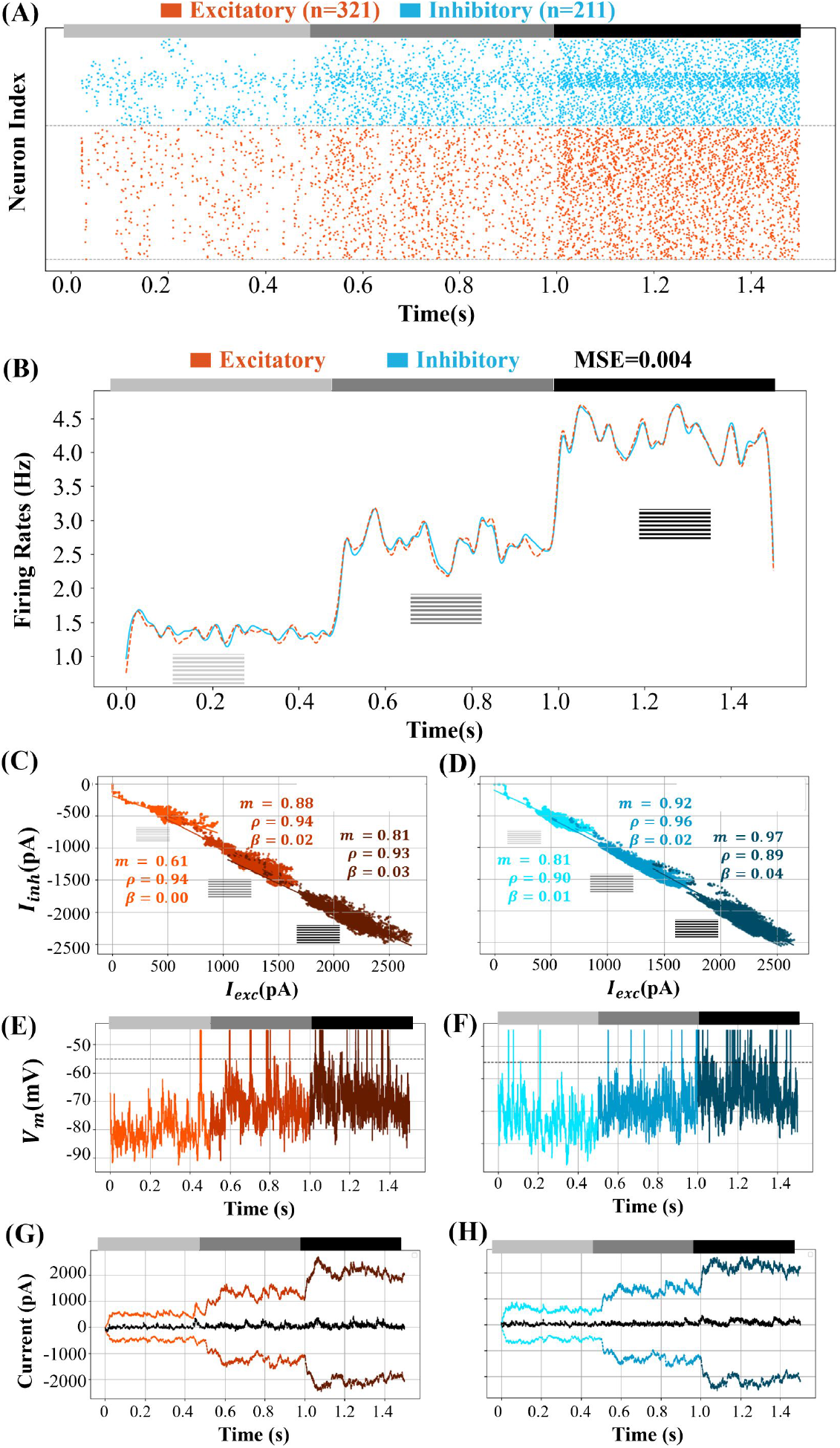
Contrast-dependent excitatory-inhibitory dynamics in response to temporally modulated Gabor stimuli. Excitatory neurons are shown in red, and inhibitory neurons are shown in blue. **(A)** Raster plot of spiking activity for excitatory (*n* = 321) and inhibitory (*n* = 211) neurons. Grayscale shading indicates contrast levels: light gray (low), gray (medium), black (high). **(B)** Population firing rates: inhibitory (solid blue) and scaled excitatory (dashed red, scaled by *a* = 4), showing tight alignment (MSE=0.004). **(C, D)** Scatter plots of excitatory (*I*_E_(*t*)) vs. inhibitory (*I*_I_(*t*)) currents for a representative excitatory (C) and inhibitory (D) neuron across contrast levels. Linear fits are segmented by contrast. **(E, F)** Membrane potential (*V*_m_(*t*)) traces for the same neurons, showing contrast-dependent depolarization and spiking. **(G, H)** Input current traces (*I*_E_(*t*), *I*_I_(*t*), *I*_n_(*t*)) for the same neurons, with standard deviations (*σ*) reported across contrast levels.

Raster plots (Fig. 11A) show increasing firing rates in both excitatory (red, *N*_E_ = 321) and inhibitory (blue, *N*_I_ = 211) populations as contrast increased. The firing rate traces (Fig. 11B) indicate that inhibitory neurons maintained higher rates overall, while the scaled excitatory population (dashed red, scaled by a factor of *a* = 4) tracked the inhibitory rate (solid blue) with a low mean squared error (MSE=0.004).

Scatter plots of synaptic currents (Fig. 11C, D) show the relationship between *I*_E_(*t*) and *I*_I_(*t*) for representative neurons under different contrast levels. Linear fits are plotted separately for each contrast segment. For the excitatory neuron (Fig. 11C), slopes *m*and correlations *ρ* increased in magnitude with contrast, while balance indices *β* decreased, indicating tighter E-I coupling at higher contrasts. Similar patterns were observed in the inhibitory neuron (Fig. 11D), suggesting that stronger visual input enhances E-I coordination.

Membrane potential traces (Fig. 11E, F) show that higher contrast levels elicited more frequent and larger depolarizations, often reaching the spiking threshold (dashed line), especially during the high-contrast epoch.

Input current traces (Fig. 11G, H) indicate that both *I*_E_(*t*) and *I*_I_(*t*) increased in magnitude with contrast, while the net current *I*_n_(*t*) = *I*_E_(*t*) + *I*_I_(*t*) remained relatively stable. The standard deviations *σ* of individual currents also increased, reflecting heightened synaptic variability under more intense stimulation. These results suggest that the network maintains E-I balance even as the drive increases, adapting dynamically to contrast changes.

### 3.6 Orientation and Feature Selectivity

To investigate the emergence of feature selectivity in the network, we presented drifting sinusoidal gratings at ten evenly spaced orientations (*θ* = 0^°^ to 180^°^ in 18^°^ increments). Each grating continuously drifted over its full spatial cycle for 2 seconds, ensuring complete coverage of spatial phases. Neurons that fired at least 10 spikes during the stimulus presentation were included in the analysis.

Feature selectivity was quantified using three complementary metrics:

- **Orientation Bias Index (OBI):** captures each neuron’s orientation selectivity across the drifting gratings, defined in Section Functional Measures and Network Readouts using a vector sum of orientation responses.
- **Phase Bias Index (PBI):** quantifies phase selectivity in an analogous manner to OBI, but based on phase-resolved responses (see Section Functional Measures and Network Readouts).
- **Modulation Ratio** (*M*_*R*_): distinguishes between simple and complex cells based on the ratio of the first harmonic (*F*_1_) to the mean (*F*_0_) of the response across phases.

Fig. 12A–B show the OBI distributions for excitatory and inhibitory populations. Orientation selectivity was prominent in excitatory neurons, with 43.6% exceeding the strong tuning threshold (OBI *>* 0.7), compared to only 3.2% of inhibitory neurons. Insets illustrate example orientation tuning and corresponding receptive fields for representative cells.

**Fig 12.**
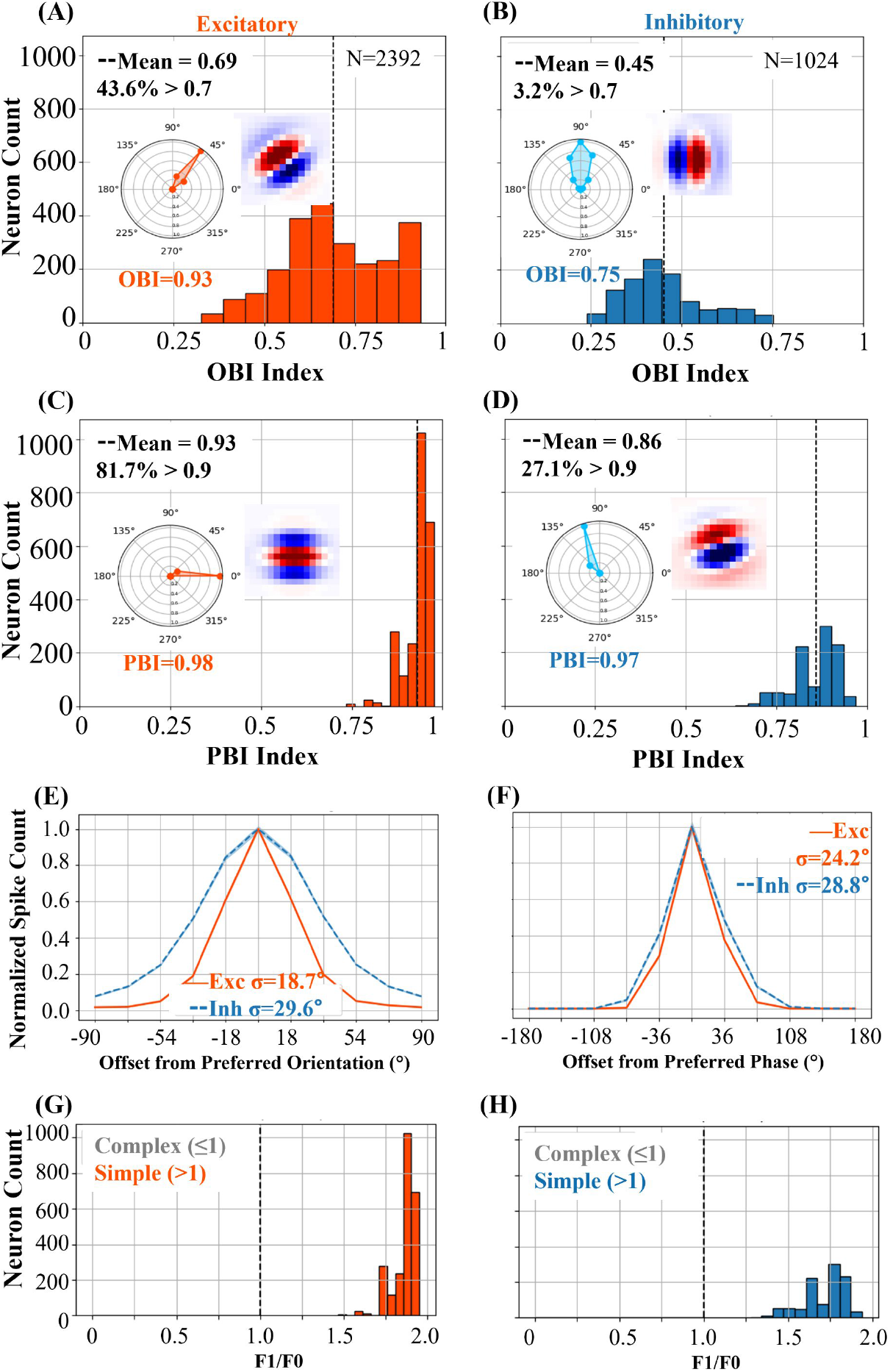
Orientation and phase selectivity in response to drifting gratings. Excitatory and inhibitory neurons in V1 layer 4 were stimulated with drifting sinusoidal gratings at 10 evenly spaced orientations (*θ* = 0^°^ to 180^°^ in 18^°^ steps), each drifting through all spatial phases. **(A–B)** Orientation Bias Index (OBI) distributions for excitatory (*n* = 2392) and inhibitory (*n* = 1024) neurons. Vertical dashed lines mark population means. Inset: example polar tuning curve and receptive field map of the most selective neuron. 43.6% of excitatory and 3.2% of inhibitory neurons exceed the OBI threshold of 0.7. **(C–D)** Phase Bias Index (PBI) distributions with a threshold at 0.9. 81.7% of excitatory and 27.1% of inhibitory neurons exceed this value. Insets show polar phase tuning and receptive fields of selected neurons. **(E–F)** Normalized and centered orientation (E) and phase (F) tuning curves. Excitatory responses are more sharply tuned, with narrower Gaussian fits: orientation tuning widths were 18.7^°^ (exc) vs. 29.6^°^ (inh), and phase tuning widths 24.2^°^ (exc) vs. 28.8^°^ (inh). **(G–H)** Distributions of modulation ratio (*F*_1_*/F*_0_), indicating that all neurons are simple-like (*F*_1_*/F*_0_ *>* 1) with most excitatory neurons clustering near the upper bound.

Phase selectivity, quantified via PBI, is presented in Fig. 12C–D. A large fraction of excitatory neurons (81.7%) surpassed the PBI threshold of 0.9, whereas only 27.1% of inhibitory neurons showed strong phase selectivity. Insets again show representative tuning and receptive fields.

Population-averaged tuning curves, normalized and aligned to each neuron’s preferred orientation or phase, are shown in Fig. 12E–F. Excitatory neurons exhibited sharper tuning than inhibitory neurons, consistent with narrower Gaussian fits: for example, the orientation tuning width (*σ*) was 18.7^°^ for excitatory and 29.6^°^ for inhibitory populations.

Finally, Fig. 12G–H shows the modulation ratio (*M*_*R*_) distributions. All excitatory and inhibitory neurons have *M*_*R*_ *>* 1, consistent with classical simple cell behavior.

Together, these findings demonstrate that drifting gratings drive strong orientation and phase selectivity in excitatory layer 4 neurons, consistent with their role in feature extraction. In contrast, inhibitory neurons display broader, less selective responses, likely reflecting their role in normalization and integration across the visual field.

### 3.7 Robustness of Network Decoding under Pixel Replacement Noise

To evaluate the robustness of the network’s decoding capability, we tested its performance under increasing levels of pixel-wise replacement noise. At each noise level, a fixed percentage of image pixels was randomly selected and replaced with values drawn from a Gaussian distribution. This simulates local corruption without altering the global image structure. The decoding process remained linear, with the reconstructed image computed as:

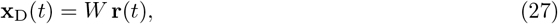

where **x**_D_(*t*) denotes the decoded (reconstructed) image at time *t, W* represents the matrix of decoding weights defined by Gabor receptive fields, and **r**(*t*) is the vector of instantaneous firing rates across the excitatory population. Each element of **x**_D_(*t*) corresponds to a pixel intensity reconstructed from the network’s population activity, providing a continuous estimate of the visual input over time.

Fig. 13A shows the Pearson correlation coefficient (*ρ*) between (i) the decoded and clean reference images (blue), (ii) the decoded output and noisy input (orange), and (iii) the noisy input and clean reference (green), across increasing noise levels.

**Fig 13.**
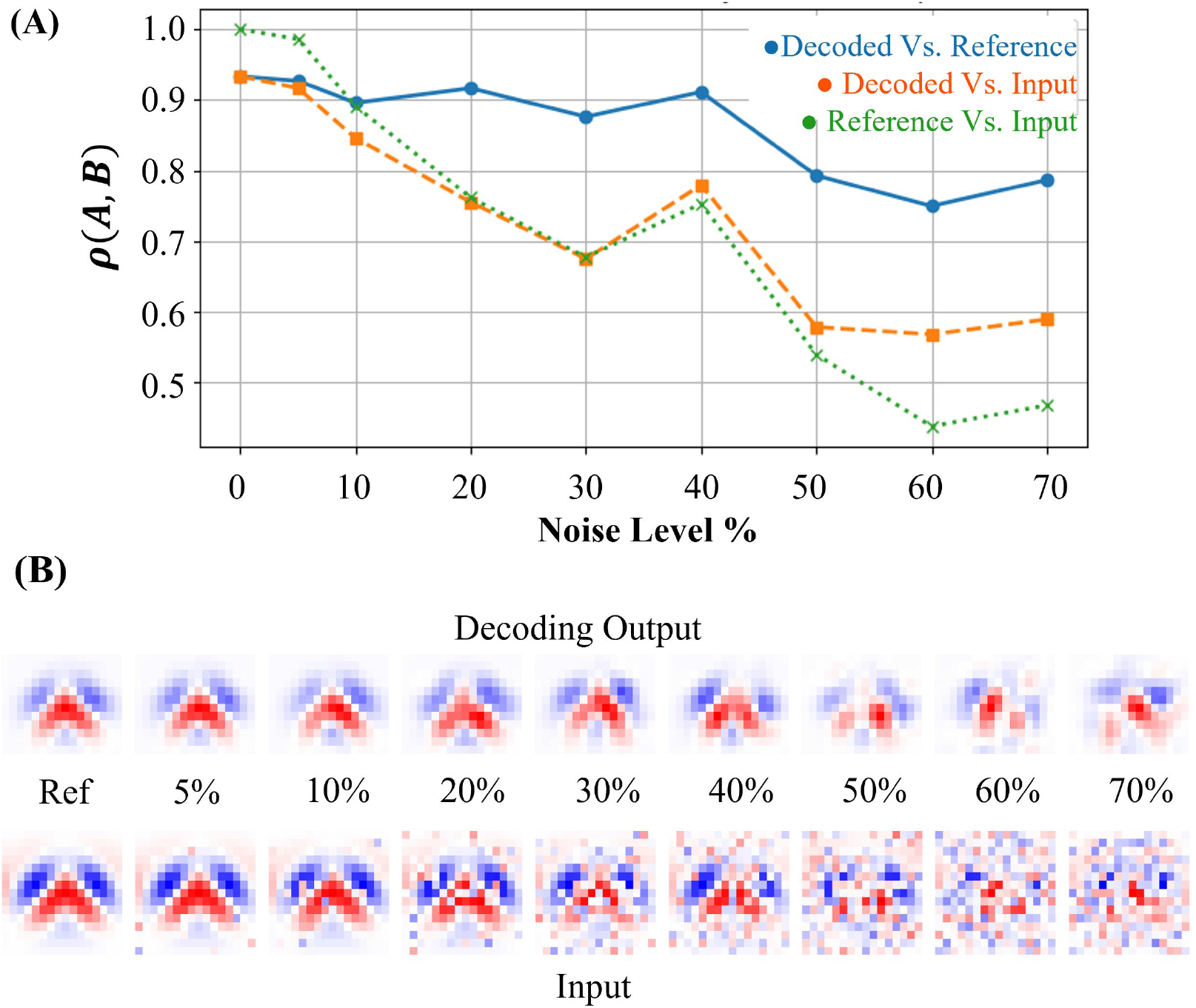
Robustness of stimulus decoding under pixel-wise noise. **(A)** Pearson correlation (*ρ*) across noise levels. Blue: decoded output vs. clean reference. Orange: decoded output vs. noisy input. Green: noisy input vs. clean reference. The decoder retains strong correlation with the reference image up to 50% pixel corruption. **(B)** Top row: Decoded outputs at increasing noise levels. Bottom row: Corresponding noisy input images, where a fixed percentage of pixels were replaced with Gaussian random values. The decoder preserves structure up to moderate noise and continues to track input patterns under severe corruption.

Visual examples in Fig. 13B (top row) show the network’s decoded outputs at each noise level. The decoder successfully reconstructed the core structure of the reference image even when up to 50% of pixels were corrupted. At higher noise levels (60–70%), reconstructions degrade but retain some coarse spatial organization.

The correlation results indicate that the decoder maintains high similarity to the reference image up to moderate noise levels. Interestingly, even as the input becomes increasingly noisy, the decoder continues to reflect the structure present in the corrupted input, as reflected by the stable correlation between the decoded output and the noisy input. This suggests that the model tracks the input structure even under severe local corruption. These results highlight the network’s capacity to robustly reconstruct meaningful visual features when a substantial portion of the input is degraded by pixel-wise noise.

## 4 Discussion

This study presented a biologically grounded spiking model of cortical layer 4, designed within the predictive coding framework. The network integrates structured thalamic input, lateral excitatory-inhibitory connectivity, and feature-specific tuning to reproduce key properties of early sensory processing in V1. By systematically analyzing responses under noisy and structured input conditions, we demonstrate how tight excitatory-inhibitory balance, sparse coding, and feature selectivity emerge from fixed connectivity alone. This section discusses the implications of these findings for predictive coding theory, biological plausibility, and future directions.

### 4.1 Predictive Coding Perspective

The model is rooted in the predictive coding framework proposed by Boerlin et. al [3], where neural spiking activity serves to iteratively minimize the mismatch between external sensory inputs and internal predictions. In this formulation, the membrane potential of each neuron encodes the instantaneous reconstruction error, while the spiking output represents an active correction of the network’s internal estimate.

Importantly, the decoding process in the model is linear: the network’s estimate of the input is reconstructed by applying fixed decoding weights to the population firing rates. Spiking occurs when the projection of a neuron’s error signal exceeds the threshold, ensuring that each spike meaningfully reduces the current mismatch.

This interpretation positions layer 4 in V1 as the bottom-most level of a hierarchical predictive coding system. Although no explicit top-down feedback is modeled here, the structured, stimulus-driven representation emerging in layer 4 provides a clean initial estimate of sensory input. In a full predictive coding hierarchy, this layer 4 output could be compared against predictions arriving at the apical dendrites of layer 2/3 neurons, enabling computation of prediction errors in deeper cortical layers.

The model thus leads to clear experimental predictions. When external input is accurately predicted, layer 4 spiking activity should decrease; conversely, when input deviates from expectation, spiking should increase to signal the mismatch. Additionally, the model predicts that inhibitory neurons should exhibit a tighter E-I balance than excitatory neurons, particularly under structured inputs such as oriented gratings. These predictions offer concrete targets for future experimental validation.

### 4.2 Biological Plausibility

Biological observations strongly constrained the model. Feedforward thalamic input was modeled using Gabor-filtered receptive fields to match simple-cell tuning properties, and a realistic 4:1 excitatory-to-inhibitory neuron ratio was maintained. Synaptic connectivity respected Dale’s Law and was fixed, without learning.

Noisy input activity reproduced realistic cortical features: low firing rates, high spike train irregularity, low pairwise synchrony, and broad membrane potential distributions. Inhibitory neurons fired more regularly and at higher rates than excitatory neurons, in line with the dynamics of fast-spiking interneurons observed in vivo.

Structured stimuli elicited reliable feature-selective responses. Both orientation and phase tuning emerged naturally from the feedforward and recurrent structure, with excitatory neurons exhibiting sharper tuning compared to inhibitory neurons. These differences were quantified using the orientation bias index (OBI) and phase bias index (PBI), as described in Section Orientation and Feature Selectivity, with their distributions shown in Fig. 12A–D. These metrics revealed sharper feature selectivity in excitatory neurons compared to inhibitory neurons, consistent with experimental findings in mouse visual cortex [59].

Furthermore, contrast modulation experiments (Section Contrast-Dependent Excitatory-Inhibitory Dynamics) revealed that the network dynamically adjusted firing rates and excitatory-inhibitory balance in response to changes in input strength. As stimulus contrast increased, spiking activity became more synchronized and depolarized, while excitatory and inhibitory currents remained tightly coordinated across contrast levels, as illustrated in Fig. 11.

Finally, decoding analyses demonstrated that the network could robustly reconstruct structured inputs, more complex than simple Gabor patterns, under moderate noise levels. Decoding fidelity, measured by Pearson’s correlation, remained high up to moderate noise amplitudes but degraded sharply under severe input corruption, consistent with known limitations in sensory robustness.

### 4.3 Contributions of This Work

This study presents a biologically grounded spiking model of cortical layer 4 that integrates predictive coding theory with realistic anatomical, physiological, and functional features of V1. The model reproduces key hallmarks of sensory processing, including irregular spiking, sparse coding, stimulus selectivity, contrast adaptation, and robust decoding, without requiring synaptic plasticity. The main contributions are grouped below:

#### Enhanced Biological Realism within the Predictive Coding Framework

- **Structured, fixed decoding weights support stable, feature-specific representations without ongoing plasticity**. Each excitatory neuron is assigned a unique Gabor-filtered receptive field based on its preferred orientation and phase (see Section Network Structure, Fig. 3A–B). These filters determine synaptic weights for feedforward input from ON/OFF LGN neurons, and also shape lateral connectivity. This biologically inspired approach enables stable, feature-selective encoding across the network without requiring plasticity mechanisms. Gabor-like receptive fields are well-established in simple cells of layer 4 in V1 [8, 9].
- **Poisson-driven LGN spike trains introduce realistic sensory variability**. Rather than using continuous-valued input as in Boerlin et al. [3], this model receives spike-based input generated from whitened and contrast-normalized natural images and gratings using a biologically grounded Poisson process (see Section Methods, Fig. 2). The ON/OFF firing rates (17 *Hz* and 8 *Hz*, respectively) are consistent with experimental observations of spontaneous LGN activity in cats and primates [44].
- **Distinct excitatory and inhibitory populations differentially contribute to feature encoding and network stabilization**. A biologically realistic 4:1 ratio of excitatory to inhibitory neurons is used (Table. 4), and all synaptic connectivity adheres to Dale’s Law. The model shows that inhibitory neurons exhibit broader, less selective tuning than excitatory neurons (see Section Orientation and Feature Selectivity, Fig. 12A–F), consistent with their integrative role observed in vivo [60, 61]. Inhibitory neurons also stabilize the network through tightly correlated and temporally matched E/I input (see Section Comparison of Excitatory-Inhibitory Balance under Noise and Grating Inputs, Fig. 10).

#### Novel Insights from Predictive Coding Simulations

- **Membrane potentials encode instantaneous reconstruction error, with spikes actively correcting the network’s internal estimate**. Consistent with the spike-based predictive coding model of Boerlin et al. [3], neurons in this model fire only when the internal estimate deviates from the sensory input, with membrane potential acting as a local error signal. This was implemented using biologically rescaled voltages and current-based dynamics (see Section Methods, especially “Biophysical Rescaling”).
- **Excitatory-inhibitory balance and sparse coding emerge without learning or plasticity**. Despite fixed weights, the model naturally exhibits tight excitatory-inhibitory balance in both spontaneous and stimulus-driven states. This is evident from the low balance index values and strong anti-correlations between *I*_E_(*t*) and *I*_I_(*t*) at both single-neuron and population levels (Section Comparison of Excitatory-Inhibitory Balance under Noise and Grating Inputs, Fig. 10). Sparse coding arises from irregular and selective spiking, particularly in the excitatory population (Section Network Response to Noise Input and Section Orientation and Feature Selectivity). These dynamics match in vivo recordings of layer 4 in cat V1 [62, 63].
- **Input structure (noise versus gratings) modulates network synchrony, spike train irregularity, and firing rate coordination**. The network transitions from irregular, asynchronous spiking under unstructured noise to more periodic and phase-locked activity under grating input, while still maintaining low pairwise correlation (Section Network Response to Grating Input, Fig. 8). This reflects known modulation of cortical state by sensory drive [64].

#### Comprehensive Characterization of Layer 4 Sensory Processing

- **Orientation and phase selectivity arise naturally from feedforward and lateral structure**. Orientation- and phase-tuned responses emerged from Gabor-structured feed-forward input and similarly tuned lateral weights. This organization led to strong selectivity in excitatory neurons without learning (see Section Orientation and Feature Selectivity, Fig. 12A–F). These properties mirror those found in layer 4 of cat and primate visual cortex [9, 64].
- **Quantitative measures (Orientation Bias Index (OBI) and Phase Bias Index (PBI)) capture distinct selectivity profiles across excitatory and inhibitory populations**. Using the OBI and PBI, the model revealed that excitatory neurons were significantly more selective than inhibitory neurons (see Section Functional Measures and Network Readouts, Section Orientation and Feature Selectivity, Fig. 12). These metrics align with those used in electrophysiological studies to characterize V1 neurons [9, 59].
- **Decoding analyses highlight the network’s robustness under moderate sensory noise and its limitations under extreme conditions**. Decoding performance, measured by reconstructing visual input from population activity, remained high under moderate pixel replacement noise (see Section Robustness of Network Decoding under Pixel Replacement Noise, Fig. 13). However, decoding fidelity sharply declined at higher noise levels, reflecting realistic constraints on sensory inference under uncertainty [65, 66].

### 4.4 Future Work

#### Laying the Foundation for Hierarchical Predictive Coding

The biologically grounded model of cortical layer 4 developed in this study forms the first stage in a hierarchical predictive coding architecture. It implements structured feedforward sensory encoding with feature selectivity, balanced excitation-inhibition, and sparse spiking, consistent with known properties of primary visual cortex.

This foundation enables the integration of higher cortical layers, particularly layer 2/3, which is modeled in the future study. Two-compartment excitatory neurons in layer 2/3 receive feedforward input from layer 4 at the soma and top-down predictions at the apical dendrite. This structure enables neurons to compute direction-specific prediction errors based on mismatches between observed input and expected feedback. This builds on predictive coding circuit models [17] and experimental studies identifying mismatch-sensitive subpopulations in layer 2/3, such as the study by Jordan and Keller [67]. The modeling of these dynamics in layer 2/3 is the focus of future research.

#### Biologically Motivated Extensions

A natural extension for future work involves incorporating synaptic plasticity into the model. This would allow decoding weights and lateral connectivity to evolve over time in response to sensory experience. Plasticity would underpin learning-dependent refinement of receptive fields, dynamic adaptation to environmental statistics, and more flexible sensory inference. Extending the study in this way would build on work proposing biologically plausible plasticity mechanisms for predictive coding in spiking networks [33, 35].

#### Model-Based Predictions for Experimental Validation

The model of layer 4 predicts specific dynamical mechanisms that can be evaluated with current experimental techniques. These predictions identify experiments that have not yet been performed but could validate the proposed computations.

##### 1. Causal influence of layer 4 excitation on prediction-error polarity in layer 2/3

The excitatory drive from layer 4 conveys bottom-up sensory evidence to the prediction-error circuit in layer 2/3, where two complementary neuronal populations encode opposite signs of mismatch. **Positive prediction-error (PE**^**+**^**) neurons** signal when sensory input is stronger than expected (*underprediction*), whereas **negative prediction-error (PE**^**–**^**) neurons** signal when sensory input is weaker than expected (*overprediction*). The model predicts that increasing layer 4 excitation amplifies PE^+^ activity and suppresses PE^−^, consistent with stronger bottom-up evidence overriding top-down predictions. Conversely, reducing layer 4 drive should release PE^−^ activity while silencing PE^+^, reflecting a state dominated by predictive feedback. Layer-specific optogenetic perturbations of excitatory neurons, combined with two-photon calcium imaging of identified PE^+^/PE^−^ populations, would directly test this predicted polarity shift in layer 2/3 driven by feedforward excitation.

##### 2. Cell-type–specific balance of excitation and inhibition

Inhibitory neurons in layer 4 are expected to maintain tighter correlations between excitatory and inhibitory currents and lower variability in E–I balance metrics than excitatory neurons, particularly under structured or predictable stimuli. These balance differences arise from the circuit’s intrinsic dynamics rather than synaptic changes. In vivo whole-cell recordings comparing E–I coupling across neuron types would validate this prediction.

##### 3. Suppression of predictable sensory drive

Layer 4 spiking represents the residual mismatch between sensory input and top-down expectations. Firing rates should decrease for well-predicted stimuli and increase for unexpected inputs, reflecting a purely dynamical suppression of redundant sensory drive. Laminar recordings during controlled prediction-violation paradigms would validate that layer 4 activity reflects dynamic precision weighting rather than static sensory drive.

##### 4. Predictability-dependent temporal sharpening

Spike latency in layer 4 neurons should shorten as stimulus sequences become more predictable, reflecting dynamic temporal refinement of feedforward signalling. Laminar multi-unit or Neuropixels recordings during temporally structured versus random visual streams would validate this untested timing effect.

##### 5. Propagation speed of prediction errors

Perturbing layer 4 input is expected to shift the timing of prediction-error propagation to layer 2/3, measurable as a transient delay in superficial calcium transients without altering magnitude. Simultaneous laminar imaging during feedforward perturbations could confirm this dynamic propagation mechanism.

Together, these experiments would establish layer 4 as the initial cortical stage that dynamically generates, times, and transmits prediction errors through stable but functionally balanced E–I interactions.

These future directions point toward a multi-layer predictive coding architecture that integrates learning, feedback, and dendritic computation to more fully capture the dynamics of cortical inference.

## 5 Acknowledgments

ANB and HM acknowledge support by the Australian Government through the Australian Research Council’s Discovery Projects funding scheme [DP220101166].

EN acknowledges support from a Melbourne Research Scholarship, and the Diane Lemaire and Dee & John Collier Travel Scholarships at the University of Melbourne.

## Supporting information

## S1 Acronym List

E-I: Excitatory-Inhibitory
ISI: Inter-Spike Interval
LGN: Lateral Geniculate Nucleus
OBI: Orientation Bias Index
PBI: Phase Bias Index
PSP: postsynaptic potential
V1: Primary Visual Cortex

## S2 Variable definitions used in the study

**Table S1.** Comprehensive variable definitions used in the V1 model.

| Variable | Description |
| --- | --- |
| Network Structure |  |
| $N_{\text{LGN,ON}}, N_{\text{LGN,OFF}}$ | Numbers of ON- and OFF-type LGN neurons |
| $N_{\text{E}}$ | Number of excitatory neurons |
| $N_I$ | Number of inhibitory neurons |
| **Gabor Parameters** |  |
| $\lambda$ | Gabor wavelength |
| $\theta$ | Gabor orientation |
| $\psi$ | Gabor phase offset |
| $\gamma$ | Gabor aspect ratio |
| $\sigma_x, \sigma_y$ | Std. dev. in $x/y$ for Gabor envelope |
| **Neuron Dynamics** |  |
| $\Delta t$ | Simulation time step |
| $\tau_m$ | Membrane time constant |
| $E_L^{(E)}, E_L^{(I)}$ | Resting potentials (E/I) |
| $R_m^{(E)}, R_m^{(I)}$ | Membrane resistances (E/I) |
| $V_r^{(E)}, V_r^{(I)}$ | Reset potentials (E/I) |
| $\tau_{r,E}, \tau_{r,I}$ | Refractory periods (E/I) |
| $\Theta_i$ | Abstract spike threshold |
| $\tilde{\Theta}_i$ | Biophysically scaled threshold |
| $a_i, b_i$ | Voltage rescaling gain and offset |
| $n_\sigma$ | Additive Gaussian noise term |
| **Synapse Dynamics** |  |
| $\tau_E, \tau_I$ | Synaptic time constants (E/I) |
| $\tau_d, \tau_s$ | Decoder and sensory time constants |
| $W_{FF}, W_{IE}, W_{EE}$ | FF, IE, and EE weight matrices |
| $W_i, \tilde{W}_i$ | Decoding vector (raw / scaled) for neuron $i$ |
| $W_{ij}^{(FF_f)}, W_{ij}^{(IE_f)}, W_{ij}^{(EE_s)}, W_{ij}^{(EI_f)}, W_{ij}^{(II_f)}$ | Pair-wise synaptic weight elements |
| **Spike Trains & Rates** |  |
| $S(t), S_i(t)$ | Generic spike train / of neuron $i$ |
| $S_j^{(LGN)}(t)$ | LGN spike train of neuron $j$ |
| $S_j^{(E)}(t), S_j^{(I)}(t)$ | Spike trains of excitatory / inhibitory neuron $j$ |
| $r(t), r_i(t)$ | Instantaneous rate / of neuron $i$ |
| $r_E(t), r_I(t), r_j^{(E)}(t)$ | Population or neuron-specific rates |
| **Currents** |  |
| $I_E(t)$ | Total excitatory current |
| $I_{E,f}^{(E)}(t), I_{I,f}^{(E)}(t), I_{E,s}^{(E)}(t)$ | Fast E, fast I, slow E currents to E population |
| $I_{E,f}^{(I)}(t), I_{I,f}^{(I)}(t)$ | Fast E/I currents to I population |
| $I_n(t)$ | Net input current |
| $c(t)$ | Decoded feedforward input current |
| **Functional Measures** |  |
| $\pi_\theta, \pi_\psi$ | Orientation / Phase Bias Indices |
| $M_R$ | Modulation ratio ($F_1/F_0$) |
| $C_V$ | Coefficient of Variation |
| $C_V^2$ | Spike-timing irregularity |
| $\rho_{H_i, H_j}$ | Spike-count cross-correlation ($i, j$) |
| $m$ | Linear slope between $I_{exc}$ and $I_{inh}$ |
| $\rho$ | Pearson correlation between excitation and inhibition |
| $\beta$ | E–I balance index |
| $\delta$ | Inter-spike interval |

